# Analysis of the influence of peptidoglycan turnover and recycling on host-pathogen interaction in the Gram-positive pathogen *Staphylococcus aureus* (Peptidoglycan recycling and Gram-positive bacteria-host interaction)

**DOI:** 10.1101/2023.07.03.547542

**Authors:** Jack Dorling, Magda Atilano, Elisabete Pires, Errin Johnson, Anna Pielach, James McCullagh, Sérgio Filipe, Petros Ligoxygakis

**Author notes:** corresponding authors: Jack Dorling and Petros Ligoxygakis.

## Abstract

During peptidoglycan recycling (PR) bacteria can recover extracellular fragments of peptidoglycan (PGN) liberated by peptidoglycan turnover (PT) during cell growth and division, and reuse them in cell wall biosynthesis or central carbon metabolism. In Gram-negative bacteria, PR has been well studied, and functions in the induction of resistance to certain classes of antibiotics, and in host-pathogen interaction. However, while Gram-negative cell envelope architecture allows for highly efficient PR, Gram-positive bacteria, which lack an outer cell membrane and are instead enclosed by a glycopolymer layer, can shed large quantities of PGN-derived material to the external environment during growth. Nonetheless, the occurrence of PR was recently demonstrated in several Gram-positive bacteria, including the Gram-positive bacterial pathogen *Staphylococcus aureus*, and its potential adaptive functions are largely unexplored. Given the known roles of PR in Gram-negative bacteria, and that Gram-positive bacteria include several important human pathogens, we asked what role PR may play during Gram-positive pathogen-host interaction. Using the model insect host *Drosophila melanogaster,* we demonstrate that *S. aureus* mutants impaired in extracellular PGN hydrolysis (Δ*atl*) and PGN fragment uptake (Δ*murP*) show differential virulence compared to their wild-type counterpart. This was linked to increased activation of the *D. melanogaster* Toll-cascade by spent supernatant from the Δ*atl* mutant. Thus, we propose that *S. aureus*, and potentially other Gram-positive bacteria, may use extracellular PGN degradation during PT to simultaneously process PGN fragments for recycling and for immune evasion, while recovery and/or metabolism of peptidoglycan fragments during PR may play more subtle roles in determining virulence.

**Author summary:** PGN is a key component of the bacterial cell wall, forming a stress-bearing sacculus surrounding the cell and providing cell shape. During growth and division, the sacculus is dynamically degraded and remodelled to ensure daughter cell separation, resulting in PT. PGN fragments released during PT can be recovered and reutilised by the cell during PR. In Gram-negative pathogens, PR is linked to antibiotic resistance, virulence and modulation of host immune recognition. In Gram-positive bacteria, PR was only recently observed. Here, we explore the roles of PT and PR in host-pathogen interaction in *S. aureus, a* Gram-positive pathogen of significant clinical relevance. Disruption of PT in *S. aureus* affected host-pathogen interaction through altering host recognition of shed PGN fragments and PR through modulation of PGN fragment recovery. This improves our understanding of the biology of this important pathogen and may aid development of novel therapeutic approaches to treat *S. aureus* infections.

## Introduction

Almost all bacteria possess a cell wall (CW) whose main structural component is the PGN sacculus [1]. PGN itself is composed of glycan strands of repeating β-1,4-linked *N*-acetylglucosamine (GlcNAc) and *N*-acetylmuramic acid (MurNAc) disaccharide aminosugar units, cross-linked by short MurNAc-linked peptides [1]. The bacterial PGN sacculus must be sufficiently rigid to resist adverse environmental conditions and rapid changes in osmotic pressure but must also be flexible enough to allow adjustment of CW shape and mechanical properties during growth, division, cell separation and differentiation. As such, the PGN sacculus is constantly remodelled during bacterial growth [2]. Remodelling is carried out by PGN hydrolases (autolysins) produced by bacteria which target covalent bonds within their own PGN sacculi [3].

PGN cleavage by autolysins can release CW-derived fragments to the surrounding environment in a process known as CW or PGN turnover. *S. aureus* exhibits CW turnover rates of ~15-25% per generation [4,5] whereas in *Escherichia coli* and *Bacillus subtilis* this is estimated at ~50% [6,7]. PR was first discovered in the Gram-negative *E. coli* [6] where diffusion of CW-derived PGN fragments (muropeptides) is restricted by the bacterium’s outer membrane, allowing efficient trapping of most turnover products and their subsequent recovery [6,8]. However, in Gram-positive bacteria such as *B. subtilis* and *S. aureus*, the lack of an outer membrane leads to shedding of large amounts of CW-derived material during growth [9,10]. Indeed, it was previously assumed that Gram-positive bacteria either do not to recycle CW material, or that the process was likely to be of little significance.

Nonetheless, it was recently discovered that Gram-positive bacteria including *S. aureus*, like Gram-negative bacteria, do indeed recycle PGN components of their CW [11,12]. In Gram-negative bacteria, the major PGN recycling substrates are GlcNAc-1,6-anhydro-MurNAc-peptide (GlcNAc-anhMurNAc) fragments [13] produced by cleavage of PGN by lytic transglycosylases which target MurNAc-β-1,4-GlcNAc bonds, generating anhMurNAc-containing muropeptides [3]. These anhydromuropeptides are then taken up via the major facilitator superfamily permease, AmpG [14]. Further catabolism by cytoplasmic PGN hydrolases produces individual aminosugars and amino acids, though larger muropeptide fragments may be directly reused [13]. Individual anhMurNAc residues are then phosphorylated by the kinase AnmK to produce *N*-acetylmuramic acid-6-phosphate (MurNAc-6-P) [15] before processing by MurQ, an etherase that converts MurNAc-6-P to GlcNAc-6-P [16].

In *E. coli*, individual MurNAc residues may also be recovered via MurP, a phosphotransferase system (PTS) component, which phosphorylates MurNAc during uptake, producing MurNAc-6-P [17]. While orthologues of AmpG are generally missing in Gram-positive bacteria, including *S. aureus*, orthologues of both MurP and MurQ are found in these bacteria [9]. Indeed, *S. aureus* can take up MurNAc from the growth medium via MurP and convert the resulting MurNAc-6-P to GlcNAc-6-P via MurQ [11]. In *S. aureus*, as in *E. coli*, recycling then proceeds via the enzyme NagA, which can deacetylate GlcNAc-6-P to produce glucosamine-6-P (GlcN-6-P) [18,19] which may then enter glycolysis after conversion to Fructose-6-phosphate (Fru-6-P), or be reused directly for PGN biosynthesis. However, unlike *E. coli*, in which recycling continues throughout growth, recycling in *S. aureus* becomes detectable only after the transition of the bacterial culture from exponential growth phase to stationary phase has begun [11,12].

*S. aureus* also extensively *O*-acetylates MurNAc residues within the PGN sacculus, rendering it extremely resistant to host-produced lysozyme-like *N*-acetylmuramidases [20]. Aside from two putative lytic transglycosylases with unknown cleavage specificity, *S. aureus* also does not appear to encode such enzymes in its genome [21]. The combined activity of peptidoglycan hydrolases of *S. aureus* is thus expected to produce MurNAc-β-1,4-GlcNAc (MurNAc-GlcNAc) PGN fragments (**Fig. 1a**), which likely represents the major PR substrate of *S. aureus*, and is taken-up via MurP in this organism [12] (**Fig. 1a**). Following uptake and concomitant phosphorylation of MurNAc-GlcNAc by MurP, MurNAc-6-P-GlcNAc is then cleaved by the cytoplasmic PGN hydrolase MupG to form MurNAc-6-P and GlcNAc [12] (**Fig. 1a**). MurNAc-6-P is then processed by MurQ as in *E. coli*. The fate of the unphosphorylated GlcNAc residue (**Fig. 1a**) is currently unknown [12]. The genes *mupG, murQ* and *murP* are encoded together in a PR operon, along with *murR*, which encodes an RpiR/AlsR family transcriptional regulator [11,22] (**Fig. 1b**).

**Figure 1.**
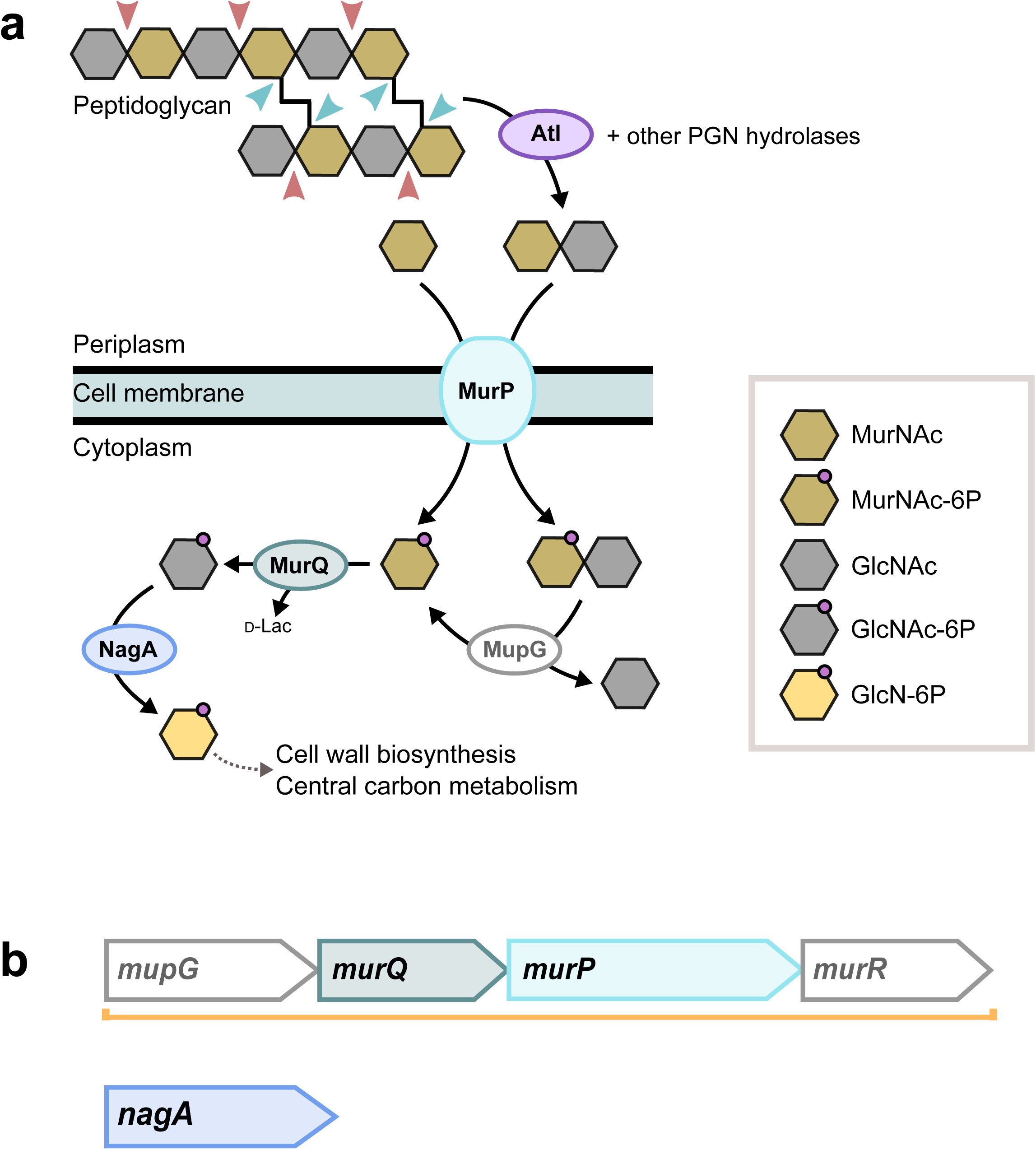
Schematic representation of peptidoglycan recycling in *S. aureus* and genomic organisation of peptidoglycan recycling genes. **a** PGN is cleaved by Atl, a bifunctional enzyme with *N*-acetylmuramoyl-L-alanine amidase (blue arrows) and *N*-acetylglucosaminidase (red arrows) activity, along with other PGN hydrolases, to produce MurNAc-GlcNAc fragments (see **Introduction**). These fragments are taken-up and phosphorylated via MurP and metabolised cytoplasmically by MupG, MurQ and NagA. The PR components studied here are shown in colour. The ‘periplasm’ is labelled following Matias *et al.* [62]. **b** The genes encoding PR genes *mupG, murQ* and *murP* are encoded in an operon along with *murR* (orange line), whereas *nagA* is not part of an operon.

Although it has now been established that PR occurs in *S. aureus* and other Gram-positive bacteria, the likely adaptive function of this process in this group is still unclear. In *B. licheniformis*, uptake of PGN-derived peptides has been implicated in the modulation of antibiotic resistance [23] while in *M. tuberculosis* antibiotic resistance induction was linked to aminosugar recycling [24]. Similarly, *S. aureus nagA* mutants are also affected in their resistance to antibiotics [19]. *S. aureus* lacking *murQ also* suffers a minor survival disadvantage during prolonged stationary phase in LB medium [11]. However, while Gram-negative PR plays roles in regulating β-lactamase expression in a number of Gram-negative species [25], it also plays roles in virulence regulation in *Salmonella enterica* serovar *Typhimurium* [26] and in regulation of host-pathogen interaction in *Neisseria* spp. and *Shigella flexneri* [27,28]. Indeed, in *M. tuberculosis,* PGN aminosugar recycling was also linked to lysozyme resistance *in vitro* [24].

Given that sugar uptake plays a major role in the pathogenic lifestyle of *S. aureus* [29] we hypothesised that PR might also play roles in host-pathogen interaction during *S. aureus* infection. To test this, we generated and characterised a panel of markerless *S. aureus* deletion mutants lacking genes encoding key components of the PR pathway in this organism, namely *murP, murQ* and *nagA* (**Fig 1a**, **b**; PR mutants), which are impaired in their ability to take up and reutilise MurNAc-containing PGN fragments, and challenged the model host *D. melanogaster* with these strains. We discovered that *S. aureus* Δ*murP,* which is unable to recover MurNAc-containing PGN fragments from the medium was compromised in its virulence in this model system, while the other two mutants, which can recover such PGN fragments but are impaired in their ability to reutilise this material, behaved as the wild-type strain.

The three mutants produced PGN and a bacterial cell surface of similar composition, and we established that the difference in their ability to kill flies or survive the innate immune system was not linked to their modified immune recognition, nor to modified lysozyme resistance as shown for other *S. aureus* mutants impaired in PGN metabolism [30]. Instead, we hypothesise that this is potentially linked to impacts on virulence regulation. In the process of conducting these experiments, we also discovered that spent culture supernatant (SCS) of *S. aureus* lacking Atl (Δ*atl*), the major autolysin of *S. aureus* (**Fig. 1a**) strongly stimulated the *D. melanogaster* immune response. This suggests that the degree of cleavage of released PGN fragments and the quantity of fragments present in the medium, which may also influence or be influenced by PR, is important in immune evasion by this organism.

## Results

### Growth parameters of PR mutants

We grew the ‘wild-type’ parental strain of *S. aureus* NCTC8325-4 (NCTC) and derived PR mutants in rich media (TSB; tryptic soy broth) to determine their growth parameters. All of the PR mutants generated in this study (**S1 Table**) showed no differences in their growth rates in rich media (**Fig. 2**, **S2 Table**; Analysis of variance (ANOVA); F_3, 8_ ^bacterial_strain^ = 3.01, p = 0.095). However, Δ*murP* and Δ*nagA* were unable to reach the same maximum OD_600_ as NCTC or Δ*murQ* (**Fig. 2**, **S2 Table**; ANOVA; F_3, 8_ ^bacterial_strain^ = 12.4, p < 0.01). Δ*murQ* also lost a smaller percentage of maximum OD_600_ after growth halted (**Fig. 2**, **S2 Table**; ANOVA; F_3, 8_ ^bacterial_strain^ = 14.5, p < 0.01). These data, in accordance with a previous report [11], demonstrate the lack of an observable impact of removal of PR enzymes during exponential growth in rich media where the bacteria are not exposed to any particular environmental stresses.

**Figure 2.**
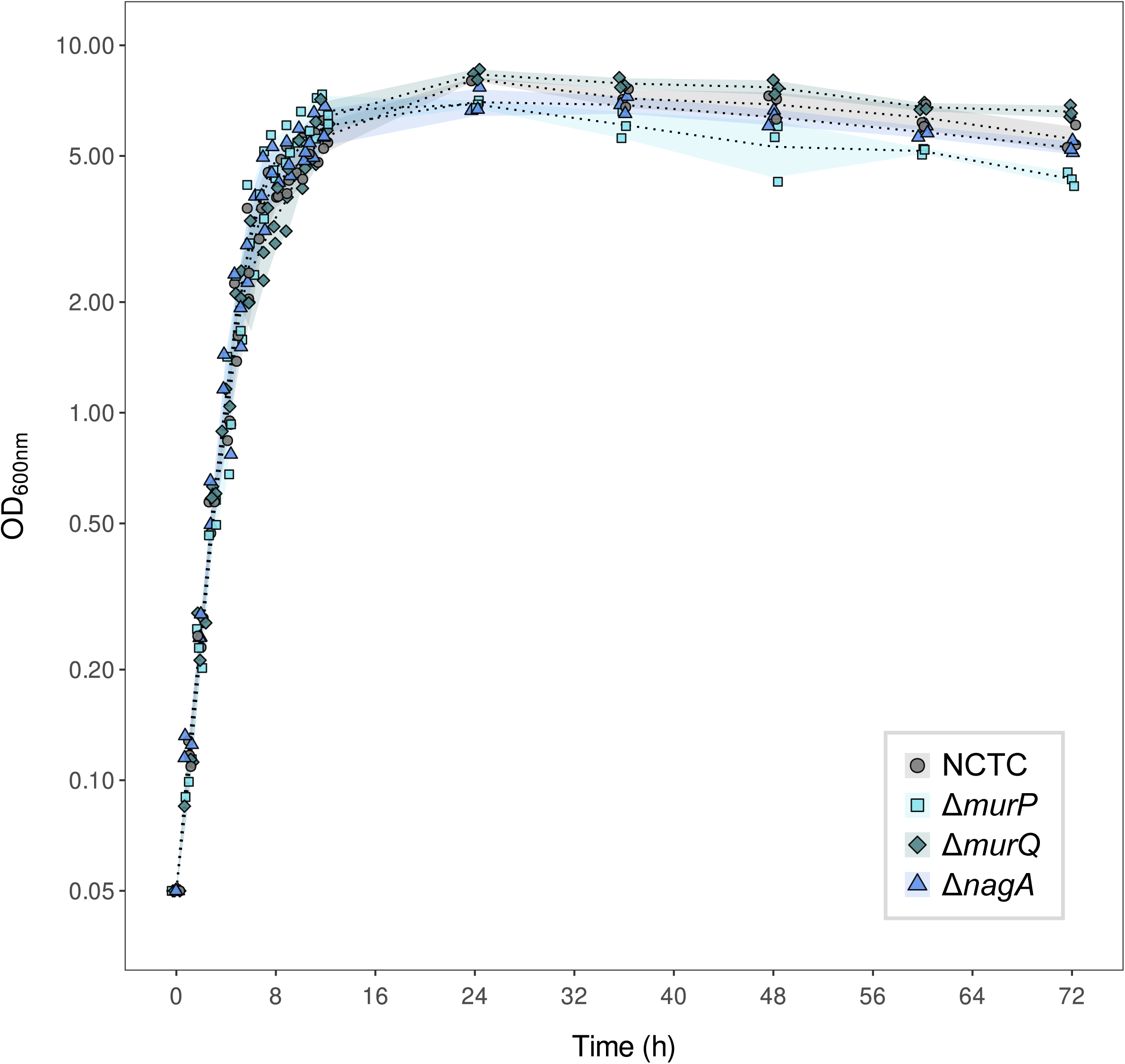
Growth of peptidoglycan recycling mutants. PR mutants were inoculated into fresh tryptic soy broth (TSB) at an initial OD_600_ of 0.05 and grown for a period of 72h. The mean OD_600_ is shown by the dotted line, and standard deviation (SD) of measurements by the shaded areas. Data are from 3 independent biological replicates. Further quantification of key growth parameters can be found in **S2 Table**.

### Dynamics of GlcNAc-6-P accumulation in NCTC and Δ*nagA*

As it has already been established that PR is most active during transition and stationary phase in *S. aureus*, and that MurNAc-6-P and MurNAc-6-P-GlcNAc accumulate in the cytoplasm during this period in mutants lacking *murQ* and *mupG*, respectively [11,12], we decided to establish whether this was also the case for GlcNAc-6-P in our Δ*nagA* mutant. In this mutant, which has a functional MurP transporter and a functional MurQ etherase capable of converting MurNAc-6-P to GlcNAc-6-P, the uptake of GlcNAc by (an)other PTS transporter(s) [28, J. Dorling unpublished data] may influence the impact of recycling of different aminosugars on *S. aureus* physiology or host-pathogen interaction. Indeed, we had already observed that Δ*nagA* reached a lower maximum OD_600_ than NCTC (**Fig. 2**).

Metabolite analysis of cytoplasmic content extracted from NCTC and Δ*nagA* grown in TSB, revealed that Δ*nagA* accumulated significantly more GlcNAc-6-P than NCTC (**Fig. 3**; Analysis of Deviance (ANODE); χ^2^ ^time_point : bacterial_strain^ = 6.97, df = 2, p < 0.001) and that this was indeed higher during transition phase (2.46 ± 0.62-fold) and stationary phase (1.51 ± 0.43-fold) than during exponential phase. Interestingly, cytoplasmic GlcNAc-6-P abundance in NCTC fell during this period (**Fig. 3**, transition; 4.81 ± 2.4-fold, stationary; 7.41 ± 7.2-fold) and in Δ*nagA* appeared to peak during transition phase (**Fig. 3**).

**Figure 3.**
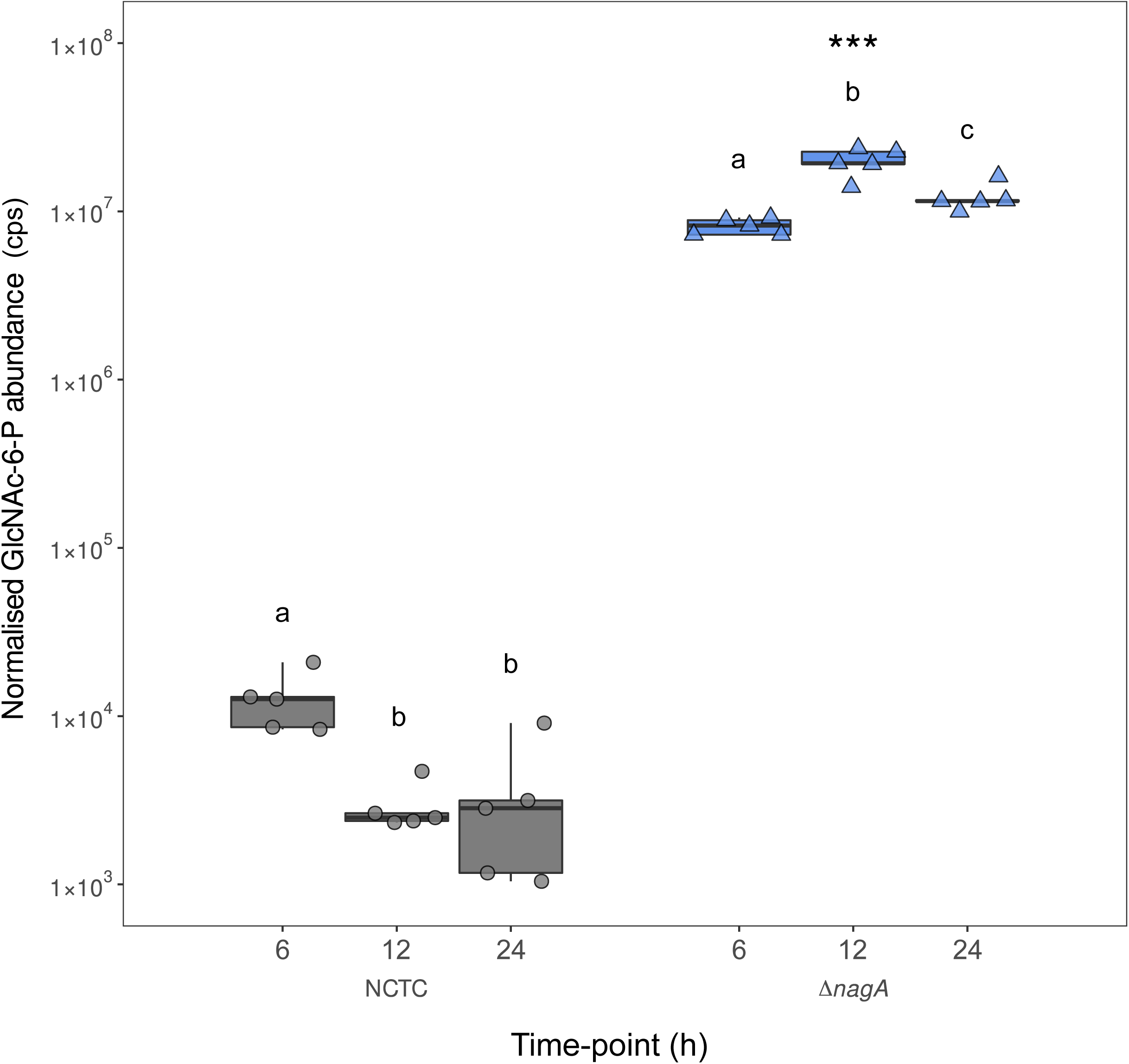
GlcNAc-6-P accumulation in Δ*nagA* throughout growth. Cytoplasmic content of bacteria grown in TSB was extracted and subjected to IC-MS/MS to quantify intracellular metabolites. GlcNAc-6-P abundance was extracted from the dataset through comparison to a reference peak generated by examination of the purified compound (see **Materials and Methods**). Data were normalised to the total abundance of all detected metabolites. cps; counts per second. Median abundance is indicated by the thick black line, while the upper and lower quartiles are given by the upper and lower limits of boxes. The upper and lower limits of the data are denoted by box whiskers. Letters given above boxes represent THSD contrasts across time points, within each strain. Samples bearing the same letter were not statistically different. Asterisks denote Tukey’s honest significant differences (THSD) post-hoc contrasts between strains; *** p < 0.001. Data are from 5 independent biological replicates.

### Impact of *nagA* deletion on downstream metabolite accumulation

NagA is the link between PR-specific metabolic activities and central carbon metabolism / PGN (re)biosynthesis [19] (**Fig. 1**). Thus, we sought to determine if the abundances of metabolites downstream of NagA were affected, to help understand whether blocking of the reutilisation of material recovered by PR may have knock-on effects on *S. aureus* metabolism, under the investigated conditions.

To do so, we examined the abundances of GlcN-6-P, the product of NagA deacetylation of GlcNAc-6-P and the hub between PR and CW biosynthesis, and Fru-6-P, the hub between PGN metabolism and glycolysis [19] (**S1 Fig**). This revealed that while no differences in the abundance of GlcN-6-P were detectable between NCTC and Δ*nagA* (**S1a Fig**; ANOVA; F_2, 26_ ^bacterial_strain^ = 0.011, p = 0.92). Fru-6-P abundances peaked in transition phase in both NCTC and Δ*nagA*, dropping in stationary phase, while still remaining at levels higher than during exponential phase (**S1b Fig**). However, the peak abundance of Fru-6-P in transition phase was lower (1.24 ± 0.21-fold) in Δ*nagA* than in NCTC (**S1b Fig**; ANOVA; F_2, 24_ ^time_point : bacterial_strain^ = 3.99, p < 0.05).

### Stationary-phase viability of PR mutants

Having now established that the dynamics of GlcNAc recycling via NagA were similar to those of MurNAc recycling, and that deletion of *nagA* led to slightly smaller pools of GlcN-6-P available for PGN biosynthesis, we then wanted to establish whether a similar minor survival disadvantage during stationary phase as that previously observed in an *S. aureus* Δ*murQ* mutant could also be observed in Δ*nagA* [11]. Thus, we tested the ability of our PR mutants to maintain viability during stationary phase in rich TSB medium (**Fig. 4**). While a slight reduction in viability of Δ*murQ* relative to other PR mutants was observed at 24h and 72h post-inoculation (ANODE; χ^2^ ^time_point : bacterial_strain^ = 4.26×10^9^, df = 12, p < 0.05), we did not document a significant reduction in Δ*murQ* viability relative to wild-type *S. aureus* (NCTC) as previously observed in LB medium [11], nor did we observe any differences in viability of the other PR mutants relative to NCTC (**Fig. 4**).

**Figure 4.**
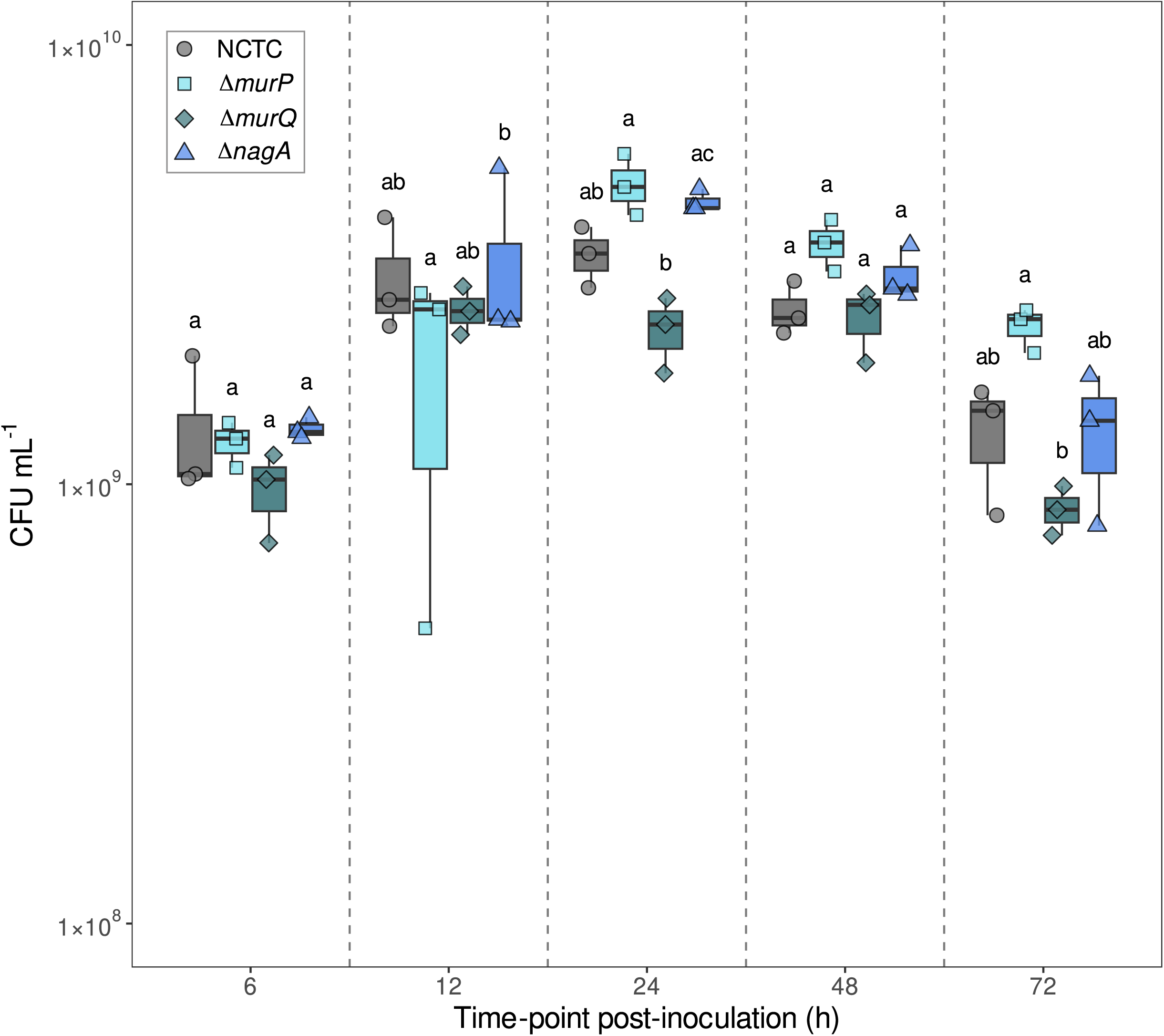
Viability of peptidoglycan recycling mutants under nutrient limitation. The number CFU present in cultures of each PR mutant at the given time points throughout growth in TSB was enumerated by plating on tryptic soy agar (TSA). The median CFU is indicated by the thick black line, while the upper and lower quartiles are given by the upper and lower limits of boxes. The upper and lower limits of the data are denoted by box whiskers. Letters given above boxes represent THSD contrasts across strains, within each time point. Samples bearing the same letter were not statistically different. Data are from 3 independent biological replicates.

### Virulence and *in vivo* bacterial load of PR mutants in *D. melanogaster*

After establishing the apparent lack of a survival disadvantage under nutrient limitation for any of our PR mutants, we wanted to address our main hypothesis; that PR may play a role in governing host-pathogen interaction in *S. aureus.* To do so, we infected *D. melanogaster* isogenic line *25174,* a line established by the *Drosophila* genetics reference panel (DGRP) [31]. We examined host survival as a proxy for bacterial virulence, as well as *in vivo* bacterial load to distinguish between the overall bacterial load within the host and their intrinsic virulence. As Atl also plays an important role in *S. aureus* PR [12], and as *S. aureus* Δ*atl* mutants are known to show reduced virulence in this model host [32], we included an *S. aureus* Δ*atl* mutant in these experiments.

These experiments revealed that while neither Δ*murQ* nor Δ*nagA* showed differential virulence when compared to NCTC, Δ*murP* was less capable of killing *D. melanogaster* over the assayed 72h period (**Fig. 5a**; log-rank test; χ^2^ ^bacterial_strain^ = 442, df = 5, p < 0.001). However, despite showing reduced virulence relative to NCTC, the impairment of virulence was far less than that of Δ*atl* (**Fig. 5a**), which showed comparable patterns of killing to those previously observed [32]. Indeed, while considerably reduced *in vivo* bacterial loads of Δ*atl* were also observed relative to NCTC (**Fig. 5b**; ANOVA; F_4, 81_ ^bacterial_strain^ = 14.2, p < 0.001), Δ*murP* had a comparable bacterial load to NCTC within the host (**Fig. 5b**).

**Figure 5.**
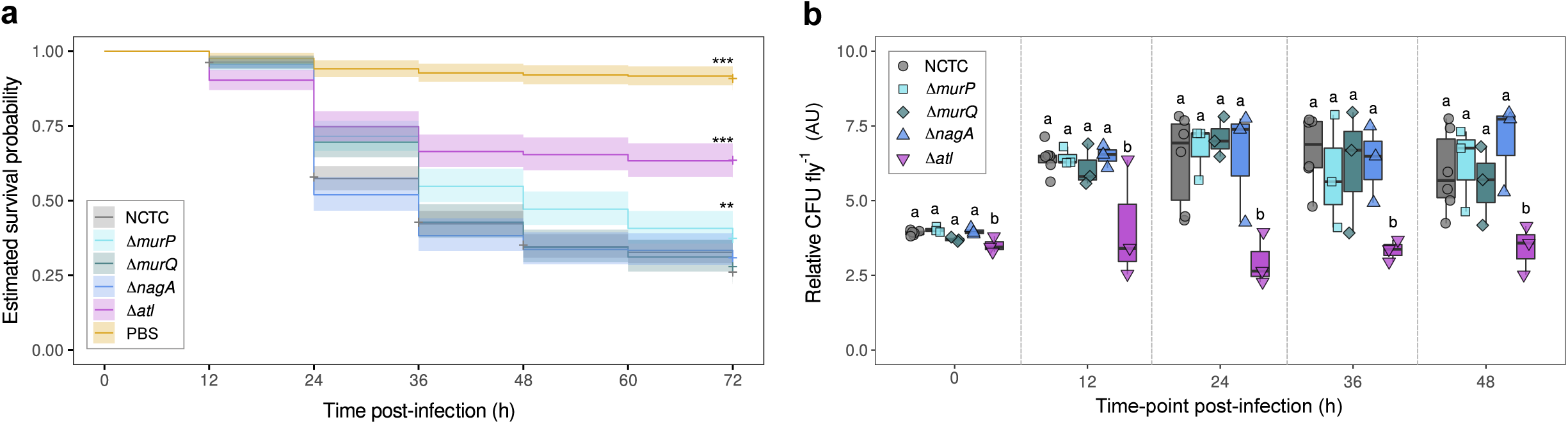
Infection of *D. melanogaster* by peptidoglycan recycling mutants. **a** *D. melanogaster 25174* flies were injected with 100-200 CFU of each of the PR mutants and *Δatl.* Their survival was monitored at 12h intervals over the course of 72h and estimated survival curves were constructed from the data. Lines represent mean estimated survival and shaded regions represent the 95% confidence intervals. Asterisks denote THSD post-hoc contrasts between strains; * p < 0.05, ** p< 0.01, *** p < 0.001. Data are from 9 independent biological replicates. Sample sizes, in number of flies injected; NCTC = 580, Δ*murP* = 312, Δ*murQ* = 299, Δ*nagA* = 306, Δ*atl* = 289, PBS = 289. **b** *D. melanogaster 25174* flies were again injected with 100-200 CFU of each of the PR mutants and *Δatl*, but this time the bacterial load (number of viable CFU fly^−1^) was enumerated every 12h for 48h. Data were box-cox transformed for analysis. AU; arbitrary units. A comparison between the original untransformed data and box-cox transformed data can be found in **S3 Table**. The median box-cox transformed bacterial load is given by the thick black line, while the upper and lower quartiles are given by the upper and lower limits of boxes. The upper and lower limits of the data are denoted by box whiskers. Letters given above boxes represent THSD contrasts across strains, within each time point. Samples bearing the same letter were not statistically different. Data are from 3 independent biological replicates, performed in 2 blocks (6 replicates for NCTC). No difference in the bacterial load was detected in NCTC between the two replicates (**S3 Fig**; ANOVA; F_1, 24_ ^experimental_block^ = 1.24, p = 0.28).

### Immune stimulation by spent PR mutant culture supernatants

Having already collected data suggesting only very subtle differences in the CW structure between PR mutants (**S2 Fig**), we reasoned that this was unlikely to explain the differences in Δ*murP* virulence we observed in **Fig. 5a**. However, as Δ*murP* mutants of *S. aureus* accumulate MurNAc-GlcNAc disaccharides extracellularly [12], which should not occur in either of the other PR mutants tested, we considered a different hypothesis. We asked whether the increased extracellular accumulation of MurNAc-GlcNAc disaccharides may potentially activate the *D. melanogaster* Toll-cascade via PGRP-SA and contribute to the reduced virulence of Δ*murP.* Multiple *S. aureus* PGN-derived molecules have been tested for their immunostimulatory activity in *D. melanogaster* [33], but no data exists for the immunostimulatory activity of this molecule, nor when present together with the infecting microorganism.

To test this hypothesis, we grew cells to stationary phase and isolated 0.22μm-filtered SCS. We then injected this into *D. melanogaster* flies containing a Drosomycin-GFP fusion (*DD1* flies), as well as their counterparts lacking PGRP-SA and the ability to detect Gram-positive PGN (*DD1^seml^* flies). We quantified GFP fluorescence in injected flies 18h post-injection (**Fig. 6**). We included Δ*atl* SCS in these experiments as a mutant expected to elicit differential immunostimulatory activity to NCTC due to reduced PGN-trimming from the cell surface of this strain [32]. In addition, any PGN fragments released from this strain may not be processed as in the parental strain as other hydrolases are present in the supernatant in differing quantities in Δ*atl* SCS [34]. We have also previously observed that polymerised muropeptides elicit a stronger immunostimulatory activity than monomeric muropeptides [33].

**Figure 6.**
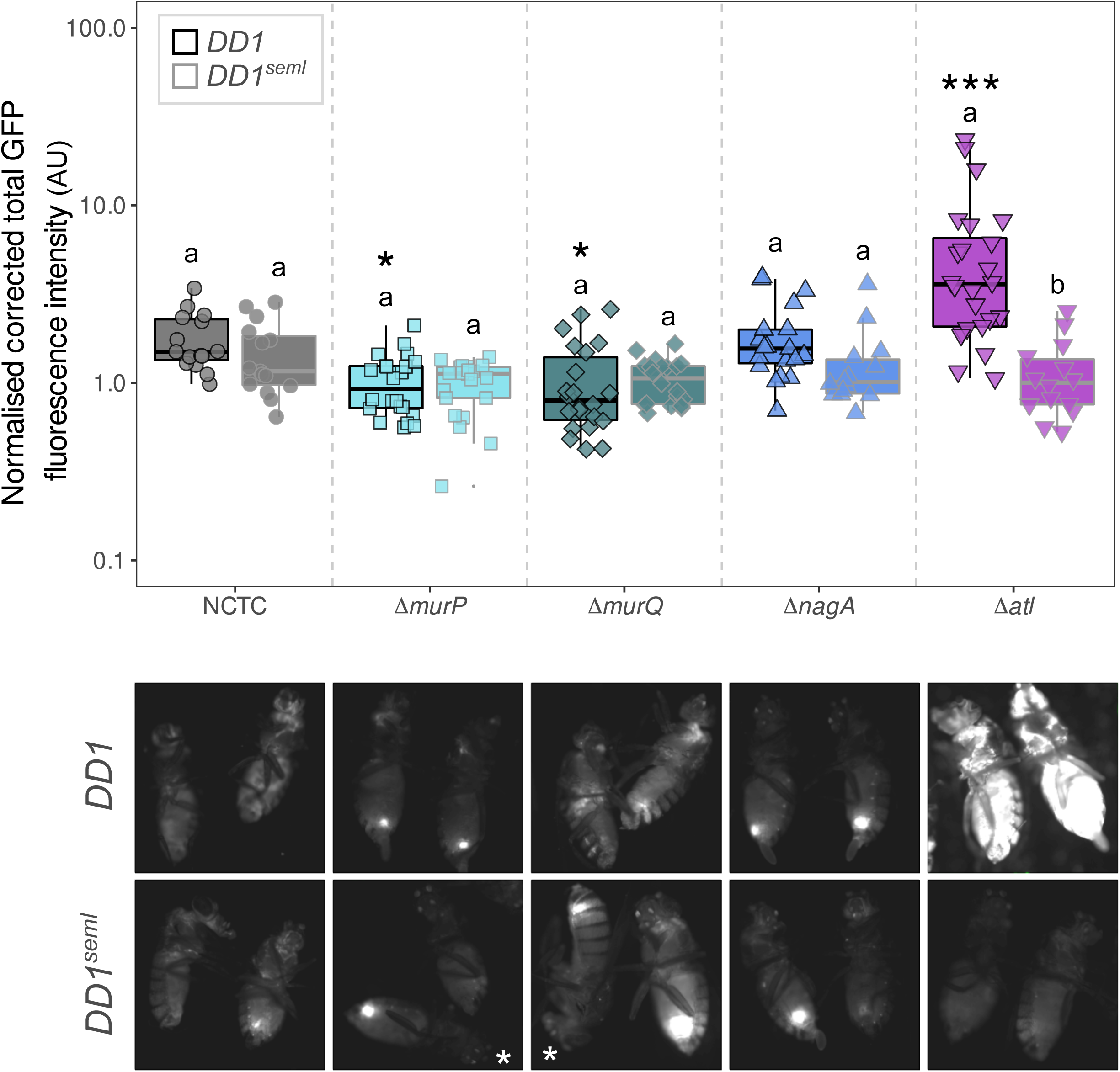
Stimulation of *D. melanogaster* immune response by spent peptidoglycan recycling mutant culture supernatant. SCS from overnight cultures of PR mutants were injected into either *DD1* (functional PGRP-SA) or *DD1^seml^* (non-functional PGRP-SA) flies. 18h later, flies were imaged to quantify Drosomycin::GFP fluorescence as a proxy for Toll-cascade activation and normalised corrected total fluorescence calculated from obtained images (see **Materials and Methods**). AU; arbitrary units. The median fluorescence is given by the thick black line, while the upper and lower quartiles are given by the upper and lower limits of boxes. The upper and lower limits of the data are denoted by box whiskers. Letters given above boxes represent THSD contrasts across fly lines, within each bacterial strain. Samples bearing the same letter were not statistically different. Asterisks denote THSD post-hoc contrasts between bacterial strains in *DD1* flies; * p < 0.05, *** p < 0.001. Data are from 3 independent biological replicates. Representative images of flies injected with SCS from each bacterial strain are shown below the plot.

We found that SCS of Δ*murQ* and Δ*murP* elicited a very modestly reduced immunostimulatory capacity in both *DD1* flies and *DD1^seml^* flies (**Fig. 6**; ANODE; χ^2^ ^bacterial_strain : fly_line^ = 47.8, df = 4, p < 0.001). However, while statistically significant, differences of such small magnitude are unlikely to be of biological significance. Unexpectedly however, we found that Δ*atl* SCS possessed a much higher immunostimulatory capacity than SCS from NCTC or the PR mutants, though only in *DD1* flies (**Fig. 6**), suggesting that this is effect is most likely linked to PGN-derived material in the SCS of Δ*atl*.

### Virulence and *in vivo* bacterial load of PR mutants in PGRP-SA-deficient *D. melanogaster* hosts

As we had not observed any differences increased immunostimulation by Δ*murP* SCS, we decided to check whether the same reduced efficiency of the killing of *D. melanogaster* by this mutant was observed in the absence of functional PGRP-SA (**Fig. 7a**). Indeed, when we infected *D. melanogaster 25714^seml^* flies, which lack a functional copy of PGRP-SA and generally die more rapidly upon infection, we observed the same reduced virulence of Δ*murP* relative to NCTC and the other PR mutants (**Fig. 7a**; log-rank test; χ^2^ ^bacterial_strain^ = 629, df = 5, p < 0.001). However, in this fly genetic background Δ*atl* showed comparable virulence to NCTC (**Fig. 7a**), as previously observed [32]. This confirmed that the differential virulence of Δ*murP* was not linked to PGRP-SA mediated recognition of PGN in this mutant. We also observed no differences in the bacterial load between any of the mutants in *25714^seml^* flies (**Fig 7b**; ANOVA; F_4, 38_ ^bacterial_strain^ = 1.57, p = 0.20).

**Figure 7.**
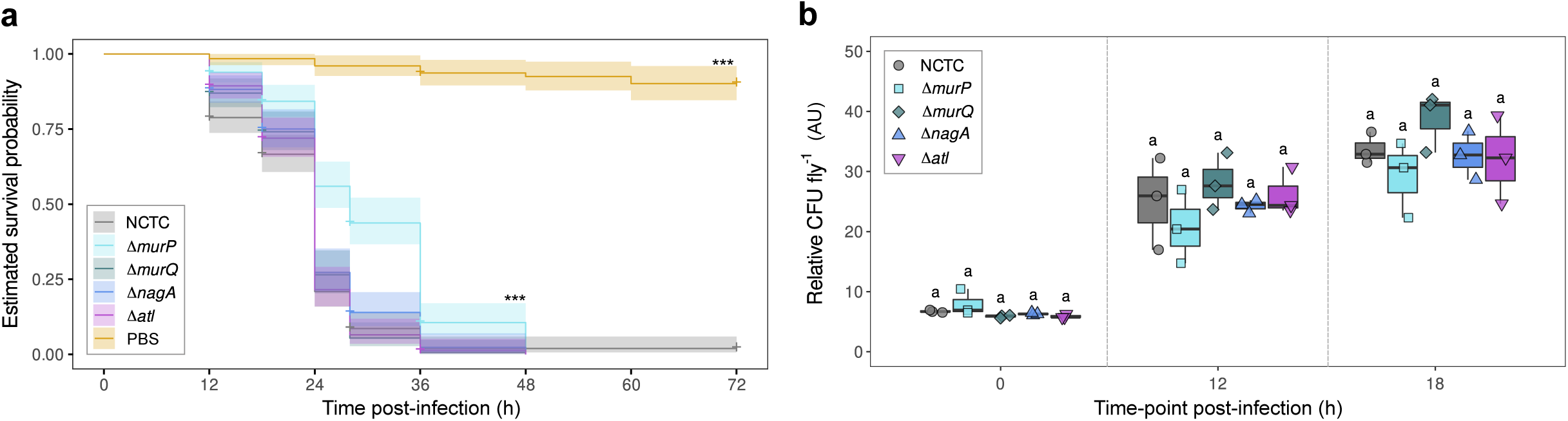
Infection of PGRP-SA deficient *D. melanogaster* by peptidoglycan recycling mutants. **a** *D. melanogaster 25174^seml^* flies were injected with 100-200 CFU of each of the PR mutants and *Δatl.* Their survival was monitored at 12h intervals over the course of 72h and estimated survival curves were constructed from the data. Lines represent mean estimated survival and shaded regions represent the 95% confidence intervals. Asterisks denote THSD post-hoc contrasts between strains; * p < 0.05, ** p< 0.01, *** p < 0.001. Data are from 7 independent biological replicates, including 3 used for determination of *in vivo* bacterial titres (**b**; see **Materials and Methods**). Sample sizes, in number of flies injected; NCTC = 236, Δ*murP* = 196, Δ*murQ* = 192, Δ*nagA* = 195, Δ*atl* = 198, PBS = 127. **b** *D. melanogaster 25174^seml^* flies were again injected with 100-200 CFU of each of the PR mutants and *Δatl*, but this time the bacterial load was measured every 12h for 48h. Data were box-cox transformed for analysis. AU; arbitrary units. A comparison between the original untransformed data and box-cox transformed data can be found in **S3 Table**. The median box-cox transformed bacterial load is given by the thick black line, while the upper and lower quartiles are given by the upper and lower limits of boxes. The upper and lower limits of the data are denoted by box whiskers. Letters given above boxes represent THSD contrasts across strains, within each time point. Samples bearing the same letter were not statistically different. Data are from 3 independent biological replicates.

### Lysozyme resistance of PR mutants

As differential virulence was not based on differential recognition of Δ*murP*, this difference in virulence had to be otherwise explained. Despite the intrinsic lysozyme resistance of *S. aureus* thanks to extensive *O*- acetylation of its PGN [20], a similar virulence phenotype, in which an *S. aureus* strain expressing a minimal PGN biosynthesis machine showed decreased virulence in both *25174* and *25174^seml^*flies, was previously explained by a decrease in lysozyme resistance in this strain [30]. Additionally, PR has also been shown to be involved in modulating lysozyme resistance via an unknown mechanism in *M. tuberculosis* [24]. We therefore reasoned that perhaps perturbation of PR might also affect lysozyme resistance. Thus, we subjected our PR mutants to a lysozyme-resistance assay (**Fig. 8**), including a Δ*tagO* mutant as a positive control known to be more sensitive to lysozyme (**S1 Table**). However, we found no difference in the lysozyme resistance of the PR mutants relative to NCTC, and instead found only an impact of *tagO* deletion (**Fig. 8**; ANOVA; F_4, 190_ ^time_point : lysozyme_treatment : bacterial_strain^ = 8.70, p < 0.001).

**Figure 8.**
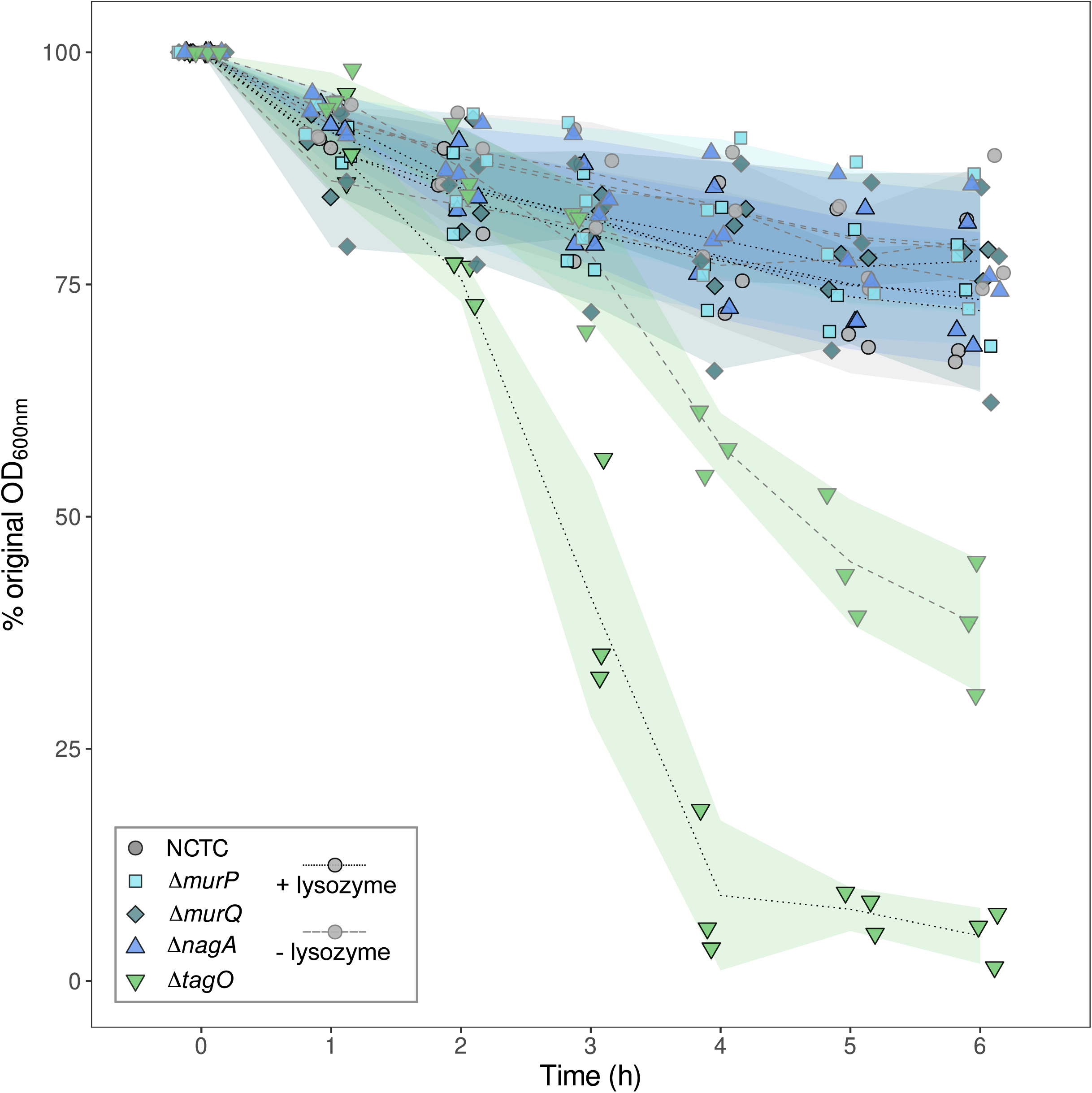
Lysozyme susceptibility of peptidoglycan recycling mutants. Overnight cultures of PR mutants and Δ*tagO* (see **S1 Table**) were washed and resuspended in PBS containing 300μg mL^−1^ lysozyme (+ lysozyme), or no lysozyme (-lysozyme). OD_600_ was then monitored over 6h. % original OD_600_ (see **Materials and Methods**) was calculated as a percentage of the OD_600_ of each strain at the start of the experiment. Mean % original OD_600_ is given by either dotted (+ lysozyme) or dashed (-lysozyme) lines and shaded areas denote SD. Data are from 3 independent biological replicates.

## Discussion

To date, only a handful of studies have addressed the topic of PR in Gram-positive bacteria [11,12,35–39] and only in recent years has PR been shown to occur in this group [11,12,24,38,40]. However, the physiological function of this process in Gram-positive bacteria remains largely unexplored with only some indication that Gram-positive PR may play a role in the maintenance of bacterial viability under nutrient limiting conditions [11,12,35] and that PR may play an important role in antibiotic and lysozyme resistance [19,23,24].

Here, while documenting some differences in growth characteristics of our PR mutants we were unable to confirm a previously detected survival defect of Δ*murQ* under nutrient-limitation [11], nor to detect such a disadvantage in our Δ*murP* or Δ*nagA* mutants. However, while both our study and that of Borisova *et al*. used rich media, we employed TSB while they used LB medium. We did however show that GlcNAc-6-P accumulated in the cytoplasm of Δ*nagA* in a similar manner to MurNAc-6-P in an *S. aureus* Δ*murQ* mutant, confirming that PR is indeed most active during stationary phase and transition phase in this organism. However, we also documented that this had very little or no impact on the downstream abundance of GlcN-6-P or Fru-6-P, and likely has little impact on PGN biosynthesis and central carbon metabolism under these conditions.

Δ*nagA* did however display a very modest reduction in Fru-6-P abundance during transition phase. GlcNAc-6-P abundance also fell in the cytoplasm of NCTC during this period. As Fru-6-P is the hub metabolite between PR and glycolysis, this suggests that PR may provide some energy for a final round of cell division during the entry into stationary phase, as suggested from studies of Gram-negative bacteria [13]. Taken together, these results suggest that while PR may function in supplying energy to the cell under nutrient limitation, its function in maintaining bacterial viability [11] is likely a minor one. However, experiments in more realistic physiological conditions, or over longer timescales, would be required to confirm this.

Here, we report that *S. aureus* mutants impaired in their ability to recover MurNAc-GlcNAc disaccharides during PR (Δ*murP*) and in the generation of these disaccharides during PGN turnover (Δ*atl*) are both less virulent than their wild-type counterparts. While the impaired virulence of Δ*atl* was already documented, we extend the characterisation of this phenotype, demonstrating that the absence of functional Atl not only increases PGRP binding to the cell surface [32], but that spent culture supernatant of Δ*atl* bacteria also elicits a robust PGRP-mediated immune response in our model host. As this was only observed in *DD1* flies possessing a functional PGRP-SA, this suggested that a decreased or aberrant hydrolysis of shed PGN-derived material was responsible for this result. While we also documented differences in the immunostimulatory capacity of SCS of Δ*murQ* and Δ*murP* these differences were small in magnitude and likely of little biological significance. We therefore could not correlate this result with the observed patterns of virulence or *in vivo* bacterial loads.

While the process of PR itself begins with the uptake of liberated PGN fragments by the cell which produced them, PGN fragments must first be generated by the action of PGN hydrolases [9]. Due to the high degree of *O*-acetylation of MurNAc residues in the PGN of *S. aureus*, this bacterium likely uses mainly, or exclusively, *N-*acetylglucosaminidases alongside amidases and endopeptidases to degrade its PGN during cell growth. *S. aureus* possesses multiple *N-*acetylglucosaminidases including Atl, the major autolysin, as well as SagA, SagB, and ScaH [41–43], which function alongside amidases and endopeptidases to generate MurNAc-GlcNAc fragments, the major PR substrate of *S. aureus* [12].

SagA, SagB, and ScaH are *N-*acetylglucosaminidases required for proper septum formation during the final stage of cell division. SagB also shortens of newly synthesized glycan strands to ensure flexibility during cell elongation [41]. Atl on the other hand is a multi-domain protein, containing *N*-terminal *N*-acetylmuramoyl-L-alanine amidase and C-terminal endo-β-*N*-acetylglucosaminidase domains [42]. Proteolytic cleavage of the Atl propeptide generates two different PGN hydrolases, which have functions in cell expansion and division, and are required for proper daughter cell separation [42,44].

Atl is already known to trim excess PGN from the bacterial cell surface, reducing PGRP binding [32]. Atl is also secreted into the external environment by *S. aureus* [42] and the two PGN hydrolases encoded by *atl* alone can generate MurNAc-GlcNac fragments. The discovery here that SCS from Atl mutants elicits a PGRP-SA dependent immune response in *D. melanogaster* highlights that Atl, and potentially other autolysins [34], also play an important role in decreasing immune stimulation by shed PGN fragments while simultaneously generating fragments that can be recycled by *S. aureus*.

PGN shedding is characteristic of many [45], but not all [39,45,46], Gram-positive bacteria, and external PGN hydrolysis is also a known feature of PR in other Gram-positives [35]. For pathogenic Gram-positive-bacteria bacteria like *S.* aureus, this may aid in avoiding immune recognition by the host, given the large quantities of PGN-derived material shed by this organism. Similarly, generation of MurNAc-GlcNAc fragments may allow cell-cell communication [47], perhaps via MurP mediated uptake of fragments originating from neighbouring *S. aureus* bacteria.

MurNAc-GlcNac fragments generated by the action of Atl and other PGN hydrolases are taken-up via MurP [12] before metabolism in the *S. aureus* cytoplasm. We also documented that Δ*murP* displayed reduced virulence when compared to its wild-type counterpart. However, unlike the reduced virulence of Δ*atl*, this phenotype could not be linked to increased recognition of accumulated of MurNAc-GlcNAc in the supernatant of this mutant [12]. Indeed, we also demonstrated that Δ*murP* displayed reduced virulence in *D. melanogaster* lacking functional PGRP-SA. It may also be possible that the increased quantities of PGN-derived fragments in the medium may activate the *D. melanogaster* immune system in a PGRP-SA independent manner, but if so this did not translate into reduced bacterial load (**Fig. 5b**, **Fig 7b**). We also established that this reduced virulence was not explained by altered lysozyme susceptibility in this mutant, as had been seen for an *S. aureus* mutant possessing minimal PGN biosynthesis machinery [30].

To try and better understand this phenotype, we turned out attention to the other genes present in the same operon as *murP* (**Fig. 1a**). One of these genes, encoding MupG, has recently been characterised and was shown to encode a cytoplasmic PGN hydrolase responsible for the cleavage of MurNAc-6-P-GlcNAc to produce MurNAc-6-P and GlcNAc [12]. The other encodes MurR [11,12] which has also been partly characterised [22]. MurR, encoded by *murR*, is also known as RpiRB and is involved in regulating pentose phosphate pathway activity and virulence factor production in *S. aureus* as a response to TCA cycle stress resulting from nutrient limitation [22]. Deletion of *murR* also results in increased production of RNAIII and a decreased rate of haemolysis [22], and therefore likely plays a role in regulation of virulence in *S. aureus*.

MurR belongs to the RpiR/AlsR family of transcriptional regulators, whose members contain highly conserved DNA-binding N-terminal helix-turn-helix domains and C-terminal sugar phosphate isomerase/sugar phosphate binding domains. The orthologue of MurR in *E. coli* [11] regulates expression of MurNAc utilisation genes in a MurNAc-6-P-dependent manner [48]. A similar interaction with MurNAc-6-P in *S. aureus* may also occur, though this is unknown. MurNAc-6-P accumulation is greatest under nutrient limitation (i.e. in stationary phase) and MurNAc-6-P may act as a signal to trigger virulence factor production via MurR. As cytoplasmic MurNAc-6-P accumulation in *murP* mutants does not occur [11,12], this could explain why the virulence of this strain is impaired. Indeed, it is becoming increasingly recognised that perturbations in metabolism alter virulence factor production and infection outcomes in *S. aureus* [49].

Accumulation of PR intermediates in the Gram-negative *Salmonella enterica* also alters virulence of this pathogen [26] and PGN metabolites are important regulatory signals involved in multiple other cellular processes in Gram-positive bacteria, including antibiotic resistance [23,47]. Alternatively, extracellularly accumulated MurNAc-GlcNAc fragments in Δ*murP* mutants [12], may bind the extracellular penicillin binding-associated and serine/threonine kinase-associated (PASTA) domain of the *S. aureus* serine-theonine kinase Stk1 [50], which is also involved in virulence regulation in this bacterium [51].

In conclusion, *S. aureus* appears to employ extracellular PGN hydrolysis to degrade fragments of PGN released as a result of cell growth and division processes to avoid activation of host immune responses, while simultaneously preparing this material for recovery by the cell. Uptake of this maximally-hydrolysed PGN-derived material [12] may then be used to support *S. aureus* metabolism to some extent, but may also influence expression of virulence, potentially via MurR-mediated virulence regulation. Ultimately, PR appears to be important in *S. aureus* host-pathogen interaction, and further investigation into the role of PR in Gram-positive bacterial virulence would be of great interest, particularly in a mammalian model host.

## Materials and Methods

### Bacterial strain construction

*S. aureus* NCTC8325-4 (NCTC) was used as the main ‘wild-type’ strain. The construction of Δ*murP*, Δ*murQ* and Δ*nagA* PR mutant strains was performed as initially described by Arnaud *et al.* [52], using the plasmids listed in **S1 Table**

To construct these mutants, we amplified ~800-900bp regions upstream (**S4 Table**; ‘p1’ and ‘p2’ primers for each respective gene) and downstream (**S4 Table**; ‘p3’ and ‘p4’ primers for each respective gene) of each respective gene. The resulting PCR products were joined by overlap PCR using ‘p1’ and ‘p4’ primers for each gene. This product was then digested with the respective restriction endonuclease enzymes (New England Biolabs) listed in **S4 Table**, allowing their subsequent ligation into a similarly digested pMAD [52] vector backbone. The constructed plasmids are listed in **S1 Table**. The plasmids were sequenced using the primers listed in **S4 Table**, and introduced into RN4220 (**S1 Table**) by electroporation. Following electroporation, plasmids were transduced using phage 80α to NCTC as previously described [53]. Insertion and excision of plasmids into the NCTC chromosome was performed as previously described [52]. Features of bacterial strains are listed in **S1 Table**.

PCR confirmation of mutant genotypes, as well as absence of the pMAD vector used for deletion, is given in **S4 Fig**. Enzymes for DNA restriction and cloning, as well as 1kB DNA ladder were purchased from New England Biolabs while GoTaq PCR reagents (Promega) were purchased from Thermo Fisher Scientific. QIAquick PCR cleanup and QIAprep Spin Miniprep kits were obtained from Qiagen. Primers were designed using Primer3plus (www.bioinformatics.nl/cgi-bin/primer3plus/primer3plus.cgi/), ReverseComplement (www.bio-informatics.org/sms/rev_comp.html) and OligoCalc (http://biotools.nubic.northwestern.edu/ OligoCalc.html) and resulting oligonucleotides purchased from Life Technologies (Thermo Fisher Scientific). Both plasmids and final deletion mutants were sequenced by Sanger sequencing to confirm the sequence of the deleted region. Primer sequences can be found in **S4 Table**.

### DNA purification

DNA was extracted from *S. aureus* for deletion fragment amplification and confirmation of mutant identity. Cells were resuspended in EDTA (50mM, pH 8.0) containing Lysostaphin and RNase A before shaking at 37°C for 30min. Further EDTA and nuclei lysis solution (Promega) were added. The mixture was incubated at 80°C for 10min and cooled to RT. Protein precipitation solution (Promega) was added and samples were vigorously mixed. Samples were incubated for 10min on ice, debris pelleted and supernatant was transferred to a fresh tube. Propan-2-ol was added mixed by inversion. Samples were centrifuged, supernatant carefully removed and samples air-dried. 70% (v/v) ethanol was added and tubes were inverted several times. Samples were centrifuged again, ethanol carefully removed and samples air-dried. DNA was dissolved in distilled water. Plasmids transformed into DH5α competent cells were purified from overnight cultures using a QIAprep Spin Miniprep Kit. DNA concentrations were measured using a Nanodrop1000 (Thermo-Fisher Scientific).

### Bacterial growth conditions

*S. aureus* strains were routinely grown in TSB (Difco) at 180rpm, or on tryptic soy agar (TSA; TSB with 1.5% added agar, Difco). Bacteria were grown at 30°C to enable comparison of results between *in vitro* and *in vivo* infection experiments (see *D. melanogaster* rearing below). Overnight cultures (~16h) were used to inoculate fresh medium at an initial optical density at 600nm (OD_600_) of 0.05. A ratio between the volumes of liquid and air of 1:5 was maintained for adequate aeration of cultures. Bacteria were plated from −80°C glycerol stocks on TSA at most 3 days before use in experiments.

### Analysis of bacterial growth parameters and viable cell counts

To analyse bacterial growth OD_600_ of bacterial cultures was measured using an Amersham Pharmacia Biochrom Ultrospec 2100 spectrophotometer. For growth experiments in TSB, r_0_ values were calculated using the `R` package `grofit` [54]. Maximum and final OD_600_ measures were extracted from the data using appropriate functions in `R` and percentage OD_600_ loss calculated as the difference between the two values divided by maximum OD_600_. For experiments examining cell viability samples were taken, placed on ice, serially diluted in fresh ice-cold TSB and 100μL of pre-determined dilutions plated with glass beads on TSA plates to achieve colony counts of ~30-300 colonies. Plates were incubated for ~30h at 30°C and photographed. Colonies were enumerated using the automatic colony counting program OpenCFU [55].

### Extraction of cytoplasmic content for metabolite analysis

Bacteria from overnight cultures were inoculated at an initial OD_600_ of 0.05 in triplicate Erlenmeyer flasks containing 200mL fresh TSB. One of each triplicate was collected at 6h, 12h and 24h of growth, OD_600_ measurements taken, and flasks chilled in an ice-ethanol bath for 10 minutes. Entire cultures were pelleted at 5000 x *g* for 15 minutes at 4°C, supernatants entirely removed by aspiration and pellets snap frozen in liquid nitrogen. Samples were stored at −80°C before further processing.

Frozen cell pellets were defrosted on ice and re-suspended to a final OD_600_ of 250. 1mL of sample was homogenised with 250mg of fine (0.25 - 0.5mm) acid-washed glass beads in a FastPrep-24 Classic (MP Biomedicals). 4 x 35s cycles of homogenisation at 6.5m s^−1^ were used, incubating samples on ice for 2 minutes after the first two cycles. Homogenised samples were pelleted at 16,000 x *g* for 10 mins at 4°C. 500μL of supernatant was then filtered through pre-washed 0.5mL 3kDa molecular weight cut-off filters (Amicon) by centrifugation at 14,000 x *g* for 20 minutes at 4°C. Filtered supernatants were then lyophilised at 55°C in a CentriVap Benchtop Centrifugal Vacuum Concentrator (Labconco) until complete dryness (~4h). Samples were then stored at −20°C.

### Metabolite profiling by IC-MS/MS and specific identification of GlcNAc-6-P

Cytoplasmic extracts were placed on ice and dissolved in 80% (v/v) LC/MS grade methanol:water. Analysis of cytoplasmic metabolite content was performed at the Mass Spectrometry Research Facility (Department of Chemistry, University of Oxford) using a Thermo Fisher Scientific ICS-5000+ ion chromatography system coupled directly to a Q-Exactive HF Hybrid Quadrupole-Orbitrap mass spectrometer with a HESI II electrospray ionisation source (Thermo Fisher Scientific), using a modified version of the previously published method [56].

A 10μL partial loop injection was used for all analyses and the chromatographic separation was performed using a Thermo Fisher Scientific Dionex IonPac AS11-HC 2×250mm ion chromatography (IC) column, (4μm particle size) with an in-line Dionex Ionpac AG11-HC 4μm 2×50mm guard column. This system incorporates an electrolytic anion generator (KOH) which produces an OH^−^ gradient from 5-100mM over 37min at a flow rate of 0.250mL min^−1^ for analyte separation. An in-line electrolytic suppressor was employed to remove OH^−^ ions and cations from the post-column eluent prior to delivery to the MS system electrospray ion source (Thermo Fisher Scientific Dionex AERS 500).

Analysis was performed in negative ion mode using a scan range of 80-900 and the resolution set to 70,000. The tune file source parameters were set as follows: sheath gas flow; 60 ms^−1^, auxiliary gas flow; 20ms^−1^, spray voltage; 3.6 V, capillary temperature; 320°C, S-lens retardation factor value; 70, heater temperature; 450°C. The automatic gain control target was set to 1×10^6^ and the maximum ionisation time value was 250ms. The column temperature was kept at 30°C throughout the experiment and full scan data were acquired in continuum mode across a mass-to-charge ratio (m/z) range of 60-900. The m/z of a GlcNAc-6-P standard (Sigma-Aldrich) was determined as 300.049 with a column retention time of 12.41 minutes (data not shown). This information was used to identify the peak of interest. Both GlcN-6-P (m/z; 258.038, retention time; 13.15 minutes) and Fru-6-P (m/z; 259.022, retention time; 14.09 minutes) were compounds already present in the compound library of the Mass Spectrometry facility at the Chemical Research Laboratory, University of Oxford. Data were acquired and analysed using Xcalibur and Progenesis software (Thermo Fisher Scientific).

### *D. melanogaster* lines, rearing and injection

*D. melanogaster* flies were raised at 25°C with a 12h:12h light:dark cycle. Flies were fed on food containing 7.69g L^−1^ agar, 34.6g L^−1^ maize, 4.15g L^−1^ soya, 7.04g L^−1^ yeast, 69.2g L^−1^ malt, and 19.2 mL L^−1^ molasses. Flies were routinely cultured in bottles containing ~50mL food, but prior to infection were housed in groups of 15-20 flies in observation vials containing ~10mL food. Fly lines used in this study are listed in **Table 1**. Flies were used 3-5 days post-eclosion as adults. Flies were shifted to 30°C 24h before infection and kept at this temperature for the duration of infection experiments. A temperature of 30°C was chosen as the survival of *D. melanogaster* is affected at 37°C, while a normal rearing temperature of 25°C for *D. melanogaster* prevents rapid bacterial growth during infection. Incubation at 30°C permits meaningful infection experiments to be carried out. This determined bacterial growth temperature for other experiments.

Overnight bacterial cultures of 20mL were pelleted at 5000 x *g* for 10 minutes at 4°C, washed twice with PBS (137mM NaCl, 2.7mM KCl, 10mM Na_2_HPO_4_, 1.8mM KH_2_PO_4_, pH 7.4) and finally re-suspended in PBS and diluted to pre-determined concentrations to ensure injection of ~100-200 CFU per fly per injection. Inoculates were prepared on ice. In the case of injection of spent bacterial culture supernatants, supernatants were saved from 20mL overnight cultures, filtered through 0.2μm filters, adjusted to an equivalent concentration of OD_600_ 5.0 and stored on ice before injection into Drosophila. Bacterial cells and spent culture supernatant samples were injected into Drosophila via the anepisternum (a soft area of the thorax, below the wing) of adult flies using a Nanoject II microinjector (Drummond Scientific) via pulled glass capillary needles.

### *D. melanogaster* survival, *in vivo* bacterial titres and immune stimulation by spent culture medium

After infection, survival of *25174* and *25174^seml^*flies was assayed at 0h, 3h, 6h, 12h and then every 4-12h for 72h. Those flies dying within the first 6h of infection were excluded from analysis as they represent casualties caused by injection. The number of flies excluded was usually between 0-2 and did not exceed 4 on any occasion. Survival data from experiments monitoring *in vivo* bacterial titres were combined with experiments used purely for assessment of survival, with censoring of flies sampled for bacterial titre determination as appropriate. In *25174^seml^* replicates 5 and 6, surviving PBS injected flies were censored at 36h as all other flies were used for CFU sampling or were dead. Bacterial infection titres were determined by collecting groups of six *25174* and *25174^seml^* flies infected in the same way as in survival experiments, starting at 0h and then every 12h for 48h. Flies were anaesthetised with CO_2_ before homogenisation in ice-cold TSB. Homogenates were kept on ice and serially diluted in fresh ice cold TSB. 100μL of pre-determined dilutions were plated by spreading on TSA to achieve colony counts of ~30-300 colonies. Platings were made in duplicate. Plates were incubated for 24-30h at 30°C and photographed. Bacterial colonies were enumerated using OpenCFU [55]. Data were collected in two blocks for *25174* flies. It was verified that NCTC counts were similar between blocks (**S3 Fig**, see also **Fig. 5b**) and data were combined for analysis.

To assess the immunostimulatory capacity of spent culture supernatants *D. melanogaster DD1* and *DD1^seml^* flies were injected with spent culture supernatant and groups of 6 flies were collected 18h after injection for imaging and assessment of GFP production. Live *D. melanogaster* flies were anaesthetised on a CO_2_ pad and imaged using an Olympus SZX-TLGAD microscope with a MVPLAPO 1X lens. Samples were illuminated using a Cool LED pE-2 coIluminator and photographed using a RETIGA R3 MONO camera. GFP signal was quantified by selecting the areas of the images occupied by flies and taking measurements of the measured area (A), the integrated density (ID) of the area (the product of the area measured and the mean grey value of that area), and mean grey value of the background (GB) before calculating CTF as follows;

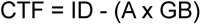

These values were then averaged over the number of flies imaged, and normalised to CTF values extracted from TSB-injected flies. Presentation images were prepared using Fiji [57]. Contrast of entire images was adjusted for presentation purposes, ensuring no clipping of high or low signals.

### Lysozyme resistance assays

Lysozyme resistance assays were carried out as in [30]. *S. aureus* cells from an overnight culture were collected by centrifugation, washed once with PBS (10mM Na_2_PO_4_, 150mM NaCl, pH 6.5), and adjusted to an OD_600_ of 0.4 in 50 ml of PBS. 20mL of suspension was placed into two 100mL flasks and incubated with or without 300 μg mL^−1^ lysozyme (final concentration; Sigma) for 6h with shaking at 30°C. Bacterial lysis was monitored by following OD_600_ and the percentage of bacterial lysis was calculated as the OD_600_ at a given time point divided by OD_600_ at 0h, multiplied by 100.

### Peptidoglycan isolation and analysis by reverse-phase high-performance liquid chromatography

PGN was prepared from exponential phase (OD_600_ 0.5-0.9) and stationary phase cells (24h post-inoculation) as previously described [33]. Briefly, cells were chilled in an ice-ethanol bath and harvested by centrifugation, resuspended in 20mL Milli-Q water and then transferred to 40mL boiling 8% (w/v) sodium dodecyl sulphate (SDS) with stirring. Samples were boiled for 30 minutes, cooled to RT and stored overnight at 4 °C. Samples were re-boiled, and SDS washed out with repeated washing with warm MilliQ water and centrifugation. SDS-free pellets were stored at −80°C.

Defrosted pellets were then homogenised with fine acid-washed glass beads in a FastPrep-24 Classic. Unbroken debris was pelleted, supernatants were retained and treated first with DNAse I and RNAse I (Sigma), then with Trypsin (Sigma). SDS was again added to a concentration of 1% (w/v) and samples boiled. Samples were washed with Milli-Q water, then resuspended in 8M LiCl for 15min at 37 °C. Samples were pelleted, resuspended in EDTA (100mM, pH 7.0) and incubated for a further 15min at 37°C, washed once more with Milli-Q water, resuspended in acetone and sonicated for 5min. Samples were washed twice more and resuspended in MilliQ water before overnight lyophilisation at 30°C. Samples were resuspended in MilliQ water to a final concentration of 20mg mL^−1^.

To remove teichoic acids, samples were treated with hydrofluoric acid (46% v/v) and incubated at 4°C for 48h. Samples were iteratively washed with Tris-HCl until the pH of the supernatant reached pH 7.0-7.5. Samples were then washed with MilliQ water twice. Samples were finally resuspended in MiliQ water, lyophilised overnight and resuspended to a final concentration of 20mg mL^−1^.

Muropeptides were prepared by digestion with mutanolysin (Sigma), reduced with sodium borohydride (Sigma) and analyzed by reverse-phase HPLC using a Hypersil ODS C-18 column (Thermo Electron Corporation) using a Shimadzu Prominence HPLC system using a 5-30% v/v methanol gradient in NaHPO_4_ at pH2.0. Sample absorbance was measured at 206nm. Data analysis was performed using Shimadzu prominence software and peaks identified where possible from comparison to previous work [58,59] and reference HPLC profiles from the Bacterial Cell Surfaces and Pathogenesis Lab (S. Filipe, ITQB, Oeiras, Portugal).

### Electron Microscopy

Bacteria from overnight cultures were inoculated into fresh TSB at an initial OD_600_ of 0.05 and grown for 24h. Cells were then collected by centrifugation, resuspended in 1mL 1% glutaraldehyde (w/v) and 1% osmium tetroxide (w/v) in 0.1M PIPES buffer on ice (0.058g L^−1^ NaCl, 0.3g L^−1^ piperazine-N,N‘bis[2-ethanesulfonic acid], 0.02g L^−1^ MgCl_2_·6H_2_O, 0.1M NaOH) and incubated at 4°C for 1h. Samples were washed with PIPES buffer and then 4 times with MilliQ water, left for 5-10 minutes between each MilliQ wash. Samples were embedded in 4% (w/v) low melting point agarose in 0.1M PIPES buffer, cut into ~1mm^3^ pieces, and incubated in 0.5% uranyl acetate overnight at 4°C in the dark. Samples were then rinsed with MilliQ water for 10min.

Samples were then serially incubated on ice in ice-cold 30%, 50%, 70%, 80%, 90% (all v/v) ethanol followed by two incubations in 100% ethanol, for 10min each. Samples were placed in anhydrous ice-cold acetone at RT for 10min. Samples were transferred to RT anhydrous acetone for another 20min. Samples were then infiltrated with low viscosity resin (TLVR; TAAB Laboratory and Microscopy equipment) by incubation in 3:1 acetone:TLVR for 1h and then 1:1 acetone:TLVR for for 2h and finally 1:3 acetone:TLVR, with rotation. Samples were incubated in TLVR overnight at RT. Resin was changed the next morning and again after another 4h.

Samples were embedded in Beem capsules filled with TLVR and resin polymerised at 60°C for 24h. Sample blocks were removed using a razor blade and ultra-thin sections made using a Diatome diamond knife using a Leica UC7 ultramicrotome and mounted on 200 mesh Cu grids. Grids were placed section-side down on a droplet of Reynolds lead-citrate and incubated at RT for 5min. Grids were washed by passing over a droplet of degassed MilliQ water, 5 times. Grids were then blotted dry and left to dry completely. Imaging was performed at 120kV using an FEI Technai 12 transmission electron microscope. Images were acquired using a Gatan OneView CMOS camera with Digital Micrograph 3.0 software.

### Statistical analyses

All statistical analyses were performed using `R` [60]. Statistical models were built including all possible interactions first (maximal models) and where appropriate (i.e. if interaction terms had little or no explanatory power) iterative model simplification was performed via likelihood ratio testing [61] with highest order non-significant interactions removed first. Non-significant interaction effects were incrementally removed and the fit of the original model and simplified model compared by ANOVA until the minimum adequate model was obtained. These models were used for analyses. General linear models were employed where possible, but where data exhibited violations of the assumptions of general linear modelling, data were transformed to conform to assumptions or generalised linear models were used instead, as most appropriate. Normality was assessed using the Shapiro-Wilk test. Error structures and link functions were chosen for generalised linear models following interpretation of diagnostics of their cognate general linear models and iterative improvement of model fitting to the data. Results are given from ANOVA tables where general linear models were used and ANODE tables for generalised linear models. ANOVA table results are presented as the F-statistic with degrees of freedom (df) in subscript and the model term in superscript, followed by the *p*-value (F-statistic_df_^model_term^ = N, p-value = n). ANODE table results are presented as the Chi-squared (χ^2^) statistic, followed by the df and the p-value (Chi-sq^model_term^ = N, df = x, p = n). Contrasts made were THSD post-hoc contrasts.

## Author contributions

Design of experiments: **JD, MLA, JM, PL, SRF**

Experimental work: **JD, MLA, EP, EJ, AP**

Analysis of data: **JD, EP**

Writing of the paper: **JD, SRF, PL**

## Supplementary Figure and Table captions

**S1 Figure. Abundance of key metabolites linking peptidoglycan recycling to peptidoglycan biosynthesis and central carbon metabolism. a** Schematic representation of the downstream metabolism of GlcNAc-6-P resulting from PR. NagB is a GlcN-6-P deaminase, converting GlcN-6-P to Fru-6-P. GlmS is an amidotransferase which converts Fru-6-P to GlcN-6-P. Cytoplasmic content of bacteria grown in TSB was extracted and subject to IC-MS/MS to quantify intracellular metabolites, from the same dataset used to create Figure 3. GlcN-6-P (**b**) and Fru-6-P (**c**) abundance was extracted from the dataset through comparison to a pre-existing compound library (see **Methods**). The corresponding symbol from **a** is given in the top-right hand corner of each plot. Data were normalised to the total abundance of all detected metabolites. cps; counts per second. The median abundance is given by the thick black line, while the upper and lower quartiles are given by the upper and lower limits of boxes. The upper and lower limits of the data are denoted by box whiskers. Letters given above box plots represent THSD contrasts across time points, within each strain. Samples bearing the same letter were not statistically different. The asterisks denote the THSD post-hoc comparison between the two strains at 12h post-inoculation; *** p < 0.001. Data are from 5 independent biological replicates.

**S2 Figure. Peptidoglycan muropeptide composition and cell wall ultrastructure of peptidoglycan recycling mutants. a** CW PGN was purified from PR mutants after either 6h (exponential phase) or 24h (stationary phase) growth. Muropeptides produced from digestion of PGN samples (see **Materials and Methods**) were then analysed by RP-HPLC and detection by UV absorption at 206nm (A_206nm_). Roman numerals I to V above the absorbance profile of NCTC for exponential phase indicate muropeptide monomers to pentamers. Peaks that differ in Δ*murP* are labelled **i** and **ii**. Peaks that differed in Δ*murQ* are shown in inset boxes and labelled **iii**. The species corresponding to peaks **i** and **ii** were identified from [58,59] and are shown as an inset. M; MurNac, G; GlcNAc. Peak **iii** was not identifiable by this method. **b** Cells from overnight cultures (Stationary phase) were fixed and images acquired by transmission electron microscopy (see **Materials and Methods**). The top row shows large fields of cells, and the lower rows high-magnification images of individual cells. Scale bars in the top row of images represent 2μm and in the lower rows 200nm.

**S3 Figure. Comparison of bacterial load over 2 experimental blocks presented in** Figure 5b. Bacterial load (number of viable CFU) per fly was enumerated every 12h for 48h. Data were box-cox transformed for analysis. AU; arbitrary units. A comparison between the original untransformed data and box-cox transformed data can be found in **S4 Table**. The median box-cox transformed bacterial load is given by the thick black line, while the upper and lower quartiles are given by the upper and lower limits of boxes. The upper and lower limits of the data are denoted by box whiskers. Each block consisted of 3 independent biological replicates.

**S4 Figure. Polymerase Chain Reaction confirmation of peptidoglycan recycling mutant construction and absence of pMAD deletion vector.** DNA was extracted from NCTC (WT; ‘wild-type’) or each of the mutants constructed in this study (**S1 Table**) and subject to PCR analysis using primers listed in **S4 Table**. **a** The absence of each of the target genes was confirmed using ‘intA’ and ‘intB’ primers. **b** The absence of the vector used for gene deletion was confirmed using primers ‘pMAD_p1’ and ‘pMAD_p2’. PCR product from PCR performed on the empty pMAD vector (pMAD) was run as a control. **c** Expected PCR product sizes for NCTC (WT) and each deletion mutant (Deletion mutant confirmation), and for each deletion vector (pMAD screening). Sizes of DNA fragments in the DNA ladder (Ladder) are given to the left of each image. Original gels from which lanes were selected are provided in **S5 Figure**.

**S5 Figure. Lanes selected from original gels to produce S4 Figure.** Orange arrows indicate the lanes which were used to produce **S4 Figure**.

## Supporting information

Dorling_et_al_Supplementary_figures_and_tables

## Acknowledgements

JD was supported by a Wellcome Trust Infection, Immunity and Translational Medicine Scholarship. We acknowledge the Sir William Dunn School of Pathology Electron Microscopy Facility for support with sample preparation and imaging.

