## Supplementary material for "Analysis of the influence of peptidoglycan turnover and recycling on host-pathogen interaction in the Gram-positive pathogen *Staphylococcus aureus* (Peptidoglycan recycling and Gram-positive bacteria-host interaction)": Dorling_et_al_Supplementary_figures_and_tables

### SUPPLEMENTARY FIGURES AND TABLES

**S1 Figure.** Abundance of key metabolites linking peptidoglycan recycling to peptidoglycan biosynthesis and central carbon metabolism.

**S2 Figure.** Peptidoglycan muropeptide composition and cell wall ultrastructure of peptidoglycan recycling mutants.

**S3 Figure.** Comparison of bacterial load over 2 experimental blocks presented in **Figure 5b**.

**S4 Figure.** Polymerase Chain Reaction confirmation of peptidoglycan recycling mutant construction and absence of pMAD deletion vector.

**S5 Figure.** Lanes selected from original gels to produce **S4 Figure**.

**S1 Table.** Bacterial strains, plasmids and fly (*D. melanogaster*) lines used in this study.

**S2 Table.** Peptidoglycan recycling mutant growth parameters.

**S3 Table.** Comparison between non-transformed and box-cox transformed CFU data for **Fig. 5b** and **7b**.

**S4 Table.** Primers used in this study.

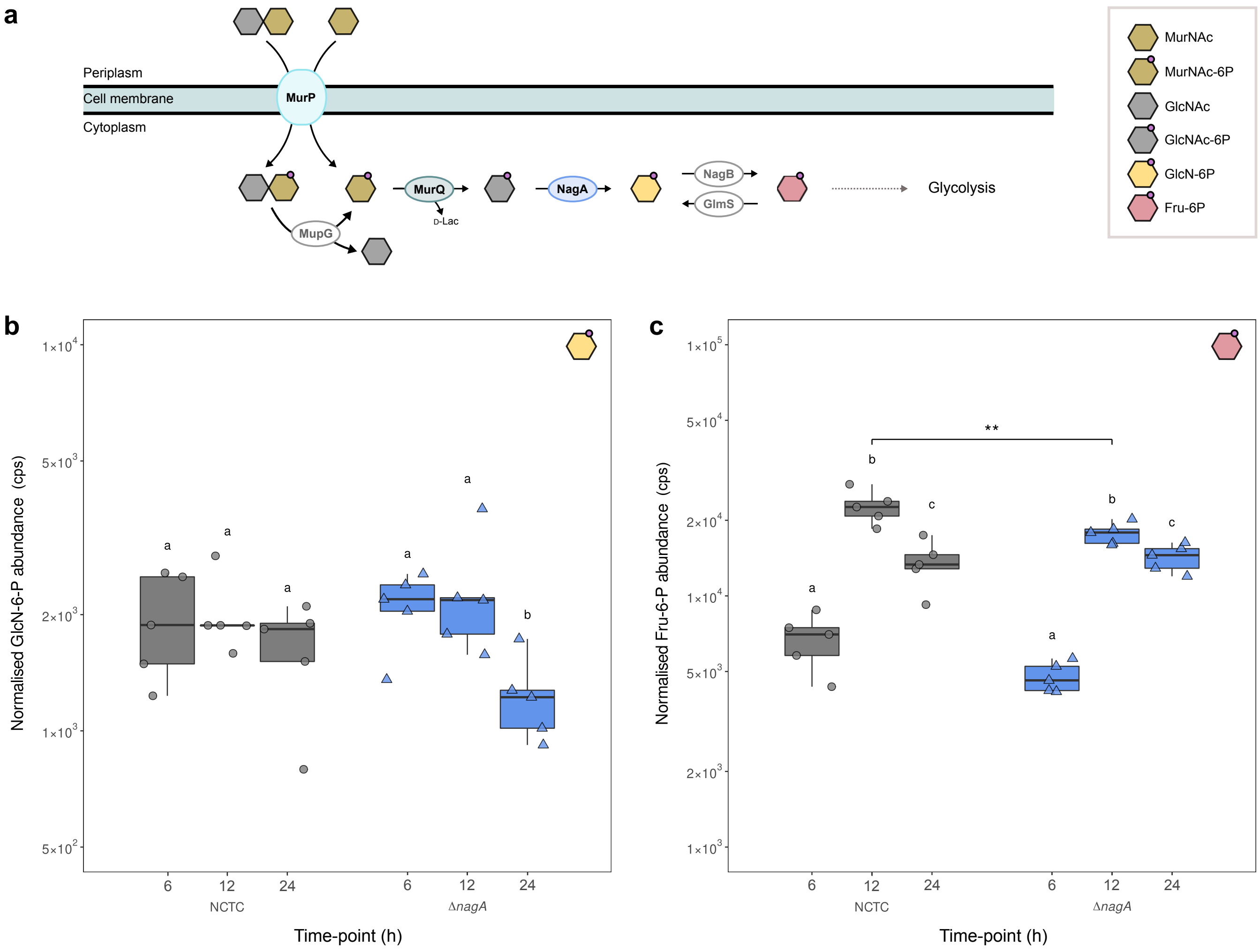
a
Periplasm
Cell membrane
Cytoplasm
MurP
MupG
MurQ
D-Lac
NagA
NagB
GlmS
Glycolysis
MurNAc
MurNAc-6P
GlcNAc
GlcNAc-6P
GlcN-6P
Fru-6P
b
Normalised GlcN-6-P abundance (cps)
1×10^4
5×10^3
2×10^3
1×10^3
5×10^2
a
a
a
a
a
b
6
12
24
6
12
24
NCTC
ΔnagA
Time-point (h)
c
Normalised Fru-6-P abundance (cps)
1×10^5
5×10^4
2×10^4
1×10^4
5×10^3
2×10^3
1×10^3
**
b
c
a
b
c
a
6
12
24
6
12
24
NCTC
ΔnagA
Time-point (h)

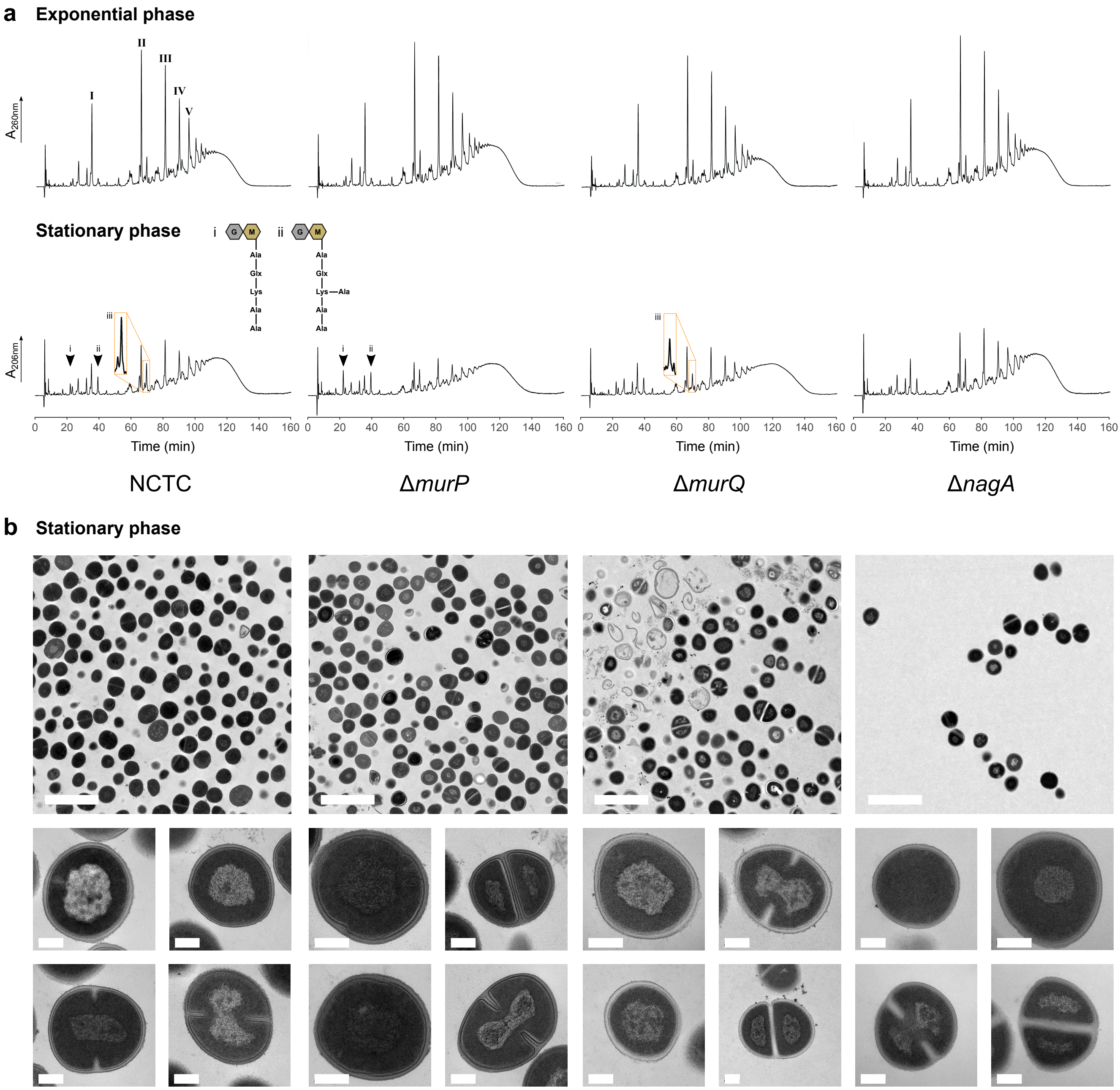

a
Exponential phase
I
II
III
IV
V
A260nm
Stationary phase
i
G
M
Ala
Glx
Lys
Ala
Ala
ii
G
M
Ala
Glx
Lys—Ala
Ala
Ala
iii
A206nm
0 20 40 60 80 100 120 140 160
Time (min)
NCTC
ΔmurP
ΔmurQ
ΔnagA
b
Stationary phase

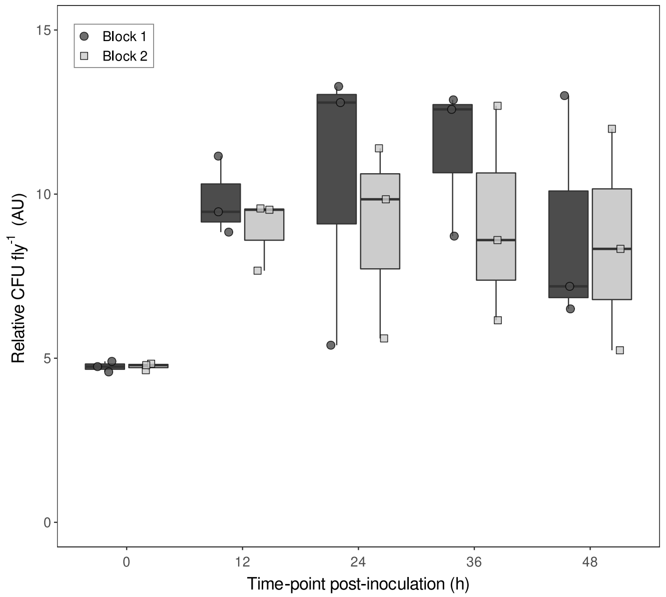
Block 1
Block 2
15
10
5
0
Relative CFU fly$^{-1}$ (AU)
0
12
24
36
48
Time-point post-inoculation (h)

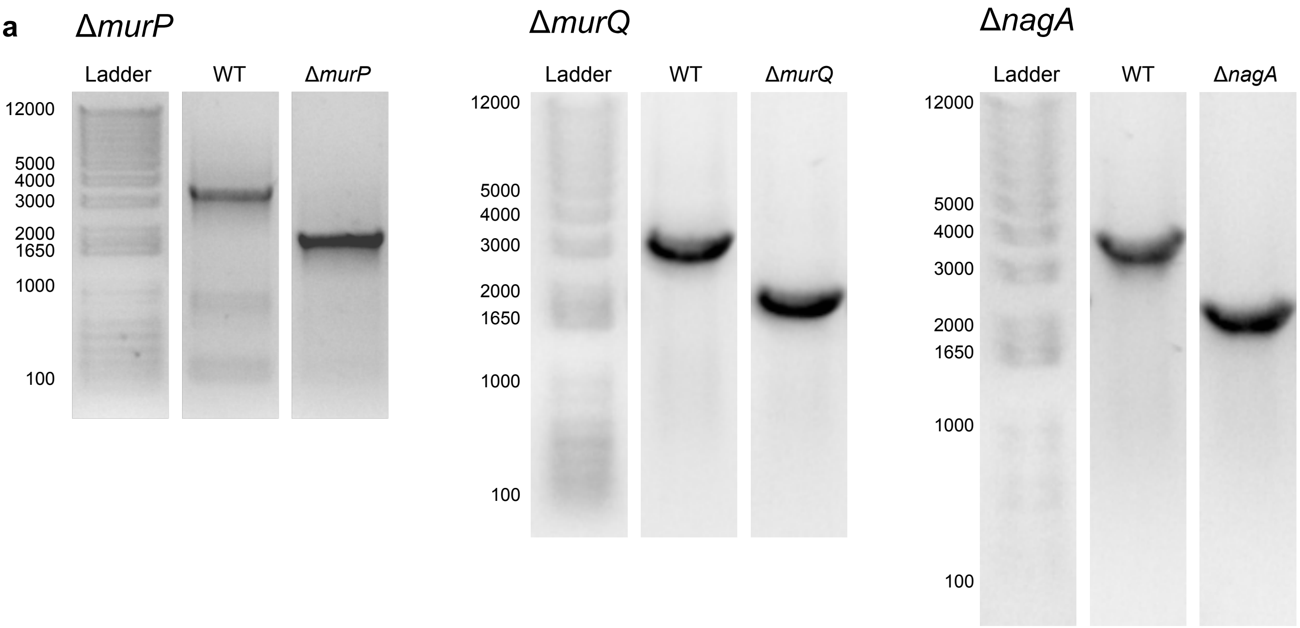


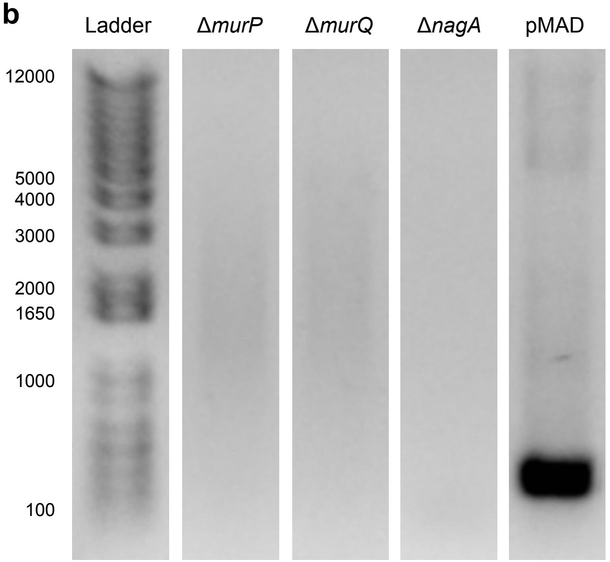


**c**

| Deletion mutant confirmation (**a**) | | |
|---|---|---|
| ***murP*** | WT: | 3330bp |
| | Δ*murP*: | 1876bp |
| ***murQ*** | WT: | 2797bp |
| | Δ*murP*: | 1902bp |
| ***nagA*** | WT: | 3290bp |
| | Δ*nagA*: | 2077bp |

| pMAD screening (**b**) | |
|---|---|
| ***murP*** | 1986bp |
| ***murQ*** | 2108bp |
| ***nagA*** | 2274bp |
| **pMAD** | 428bp |

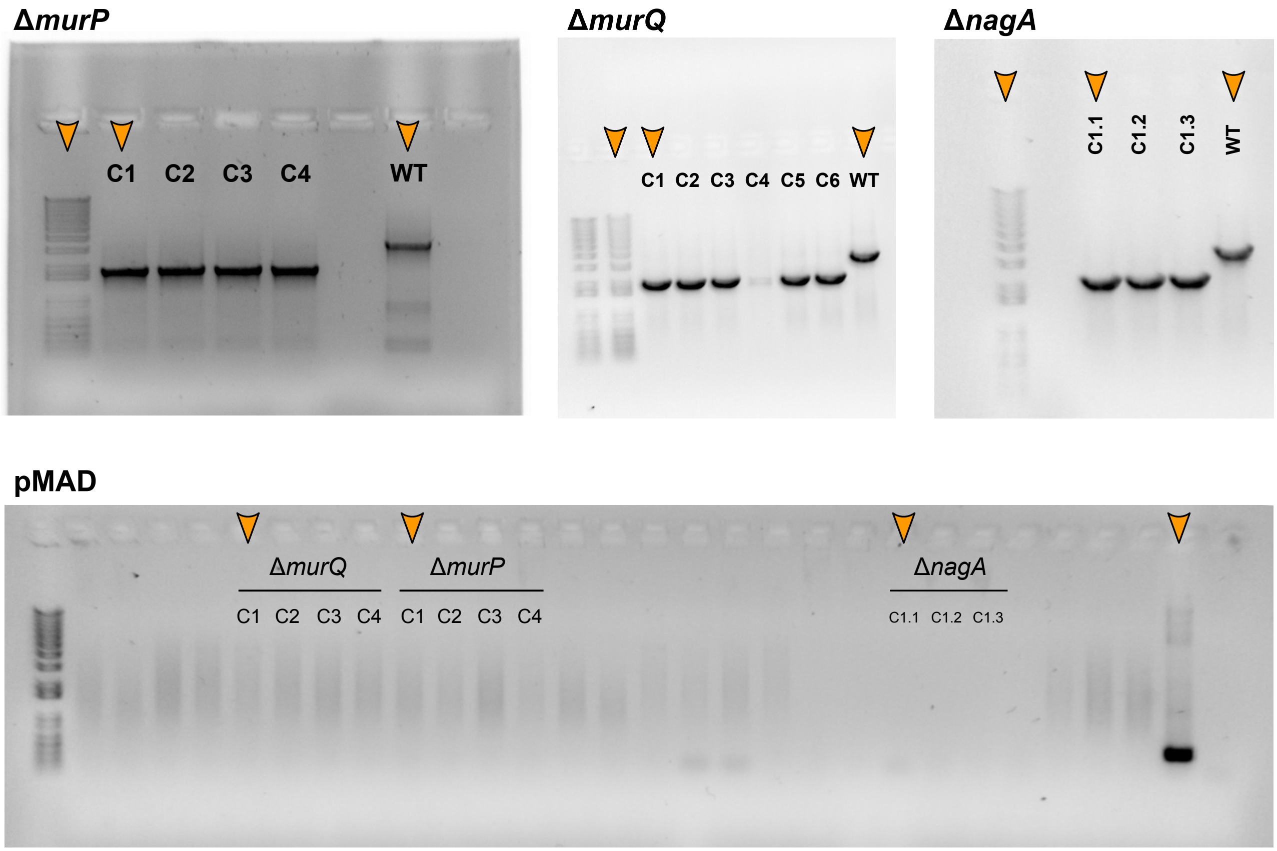
ΔmurP
C1
C2
C3
C4
WT
ΔmurQ
C1
C2
C3
C4
C5
C6
WT
ΔnagA
C1.1
C1.2
C1.3
WT
pMAD
ΔmurQ
C1
C2
C3
C4
ΔmurP
C1
C2
C3
C4
ΔnagA
C1.1
C1.2
C1.3

**S1 Table.** Bacterial strains, plasmids and fly (*D. melanogaster*) lines used in this study.

| Strain / Plasmid / Fly line | Characteristics | Source |
|---|---|---|
| ***Bacterial strains*** | | |
| *E. coli* DH5α | *E. coli* cloning strain | Lab stock |
| RN4220 | Restriction-deficient derivative of NCTC8325-4 with an inactivated restriction system permitting its use as an intermediate cloning strain | R. Novick |
| NCTC | *S. aureus* reference strain, NCTC8325 derivative lacking prophages Φ11, 12 and 13. | R. Novick |
| Δ*murP* | NCTC8325-4 *murP* null mutant | This study |
| Δ*murQ* | NCTC8325-4 *murQ* null mutant | This study |
| Δ*nagA* | NCTC8325-4 *nagA* null mutant | This study |
| Δ*atl* | NCTC8325-4 *atl* null mutant | [1] |
| Δ*tagO* | NCTC8325-4 *tag* null mutant | [2] |
| ***Plasmids*** | | |
| pMAD | *E. coli – S. aureus* (Gram-positive) shuttle vector with a thermosensitive origin of replication in Gram-positive bacteria; $Amp^R$ $Ery^R$ $LacZ^+$ | [3] |
| pMADΔ*murP* | pMAD + *murP* deletion fragment | This study |
| pMADΔ*murQ* | pMAD + *murQ* deletion fragment | This study |
| pMADΔ*nagA* | pMAD + *nagA* deletion fragment | This study |
| ***Fly lines*** | | |
| *25174* | DGRP-208. Drosophila Genetic Reference Panel (DGRP) strain - sequenced isogenic reference wild type strain. | BDSC [1] |
| $25174^{seml}$ | *25174* carrying a C80Y mutation in PGRP-SA | M. L. Atilano |
| *DD1* | *y w drs-GFP dpt-LacZ* reference line | BDSC |
| $DD1^{seml}$ | *DD1*carrying a C80Y mutation in PGRP-SA | [4] |

**Abbreviations;** Amp; ampicillin, Ery; erythromycin, Bloomington Drosophila stock centre, *y*; yellow body, *w*; white eyes

**S2 Table.** Peptidoglycan recycling mutant growth parameters.

| ***S. aureus* strain** | **Mean growth rate ± SD** | **Mean max. $OD_{600}$ ± SD** | **Mean final $OD_{600}$ ± SD** | **Mean % loss in $OD_{600}$ ± SD** |
|---|---|---|---|---|
| NCTC | 0.293 ± 0.067 [a] | 8.08 ± 0.09 [a] | 5.57 ± 0.44 [a] | 31.1 ± 5.5 [a] |
| Δ*murP* | 0.380 ± 0.053 [a] | 7.00 ± 0.31 [b] | 4.32 ± 0.19 [b] | 38.2 ± 0.73 [a] |
| Δ*murQ* | 0.241 ± 0.052 [a] | 8.34 ± 0.27 [a] | 6.59 ± 0.25 [c] | 20.9 ± 2.5 [b] |
| Δ*nagA* | 0.328 ± 0.060 [a] | 7.08 ± 0.53 [b] | 5.27 ± 0.21 [a] | 25.5 ± 3.1 [a] |

Letters given beside values in each column represent THSD contrasts across bacterial strains. Samples bearing the same letter were not statistically different.

**S3 Table.** Comparison between non-transformed and box-cox transformed CFU data for **Fig. 5b** and **7b**.

| Fly line | Time-point | Bacterial strain | Median untransformed CFU fly$^{-1}$ [A] | Mean box-cox transformed relative CFU fly$^{-1}$ (AU) [B] |
|---|---|---|---|---|
| **Fig. 5b** | | | | |
| *25174* | 0 | NCTC | $1.50 \times 10^{2}$ | 3.92 |
| | | Δ*murP* | $1.68 \times 10^{2}$ | 4.02 |
| | | Δ*murQ* | $1.01 \times 10^{2}$ | 3.70 |
| | | Δ*nagA* | $1.50 \times 10^{2}$ | 3.97 |
| | | Δ*atl* | $6.92 \times 10^{1}$ | 3.51 |
| *25174* | 12 | NCTC | $3.79 \times 10^{4}$ | 6.41 |
| | | Δ*murP* | $2.13 \times 10^{4}$ | 6.45 |
| | | Δ*murQ* | $6.33 \times 10^{3}$ | 6.10 |
| | | Δ*nagA* | $4.45 \times 10^{4}$ | 6.49 |
| | | Δ*atl* | $2.58 \times 10^{1}$ | 2.89 |
| *25174* | 24 | NCTC | $2.40 \times 10^{5}$ | 6.36 |
| | | Δ*murP* | $4.60 \times 10^{5}$ | 6.73 |
| | | Δ*murQ* | $1.85 \times 10^{5}$ | 7.09 |
| | | Δ*nagA* | $7.67 \times 10^{5}$ | 6.47 |
| | | Δ*atl* | $2.17 \times 10^{1}$ | 2.96 |
| *25174* | 36 | NCTC | $1.02 \times 10^{6}$ | 6.67 |
| | | Δ*murP* | $4.15 \times 10^{3}$ | 5.87 |
| | | Δ*murQ* | $6.83 \times 10^{4}$ | 6.19 |
| | | Δ*nagA* | $3.75 \times 10^{4}$ | 6.30 |
| | | Δ*atl* | $6.17 \times 10^{1}$ | 3.34 |
| *25174* | 48 | NCTC | $5.77 \times 10^{3}$ | 5.96 |
| | | Δ*murP* | $8.50 \times 10^{4}$ | 6.23 |
| | | Δ*murQ* | $4.83 \times 10^{3}$ | 5.56 |
| | | Δ*nagA* | $3.33 \times 10^{6}$ | 6.98 |
| | | Δ*atl* | $8.50 \times 10^{1}$ | 3.42 |
| **Fig. 7b** | | | | |
| $25174^{seml}$ | 0 | NCTC | $1.65 \times 10^{2}$ | 6.72 |
| | | Δ*murP* | $1.92 \times 10^{2}$ | 7.95 |
| | | Δ*murQ* | $1.07 \times 10^{2}$ | 5.88 |
| | | Δ*nagA* | $1.30 \times 10^{2}$ | 6.30 |
| | | Δ*atl* | $9.33 \times 10^{1}$ | 5.92 |
| $25174^{seml}$ | 12 | NCTC | $3.40 \times 10^{5}$ | 25.05 |
| | | Δ*murP* | $6.50 \times 10^{4}$ | 20.72 |
| | | Δ*murQ* | $5.35 \times 10^{5}$ | 28.14 |
| | | Δ*nagA* | $2.27 \times 10^{5}$ | 24.25 |
| | | Δ*atl* | $2.18 \times 10^{5}$ | 26.21 |
| $25174^{seml}$ | 18 | NCTC | $1.97 \times 10^{6}$ | 33.66 |
| | | Δ*murP* | $1.15 \times 10^{6}$ | 29.21 |
| | | Δ*murQ* | $1.11 \times 10^{7}$ | 38.75 |
| | | Δ*nagA* | $1.90 \times 10^{6}$ | 32.70 |
| | | Δ*atl* | $1.70 \times 10^{6}$ | 32.07 |

**S4 Table.** Primers used in this study.

| Primer | Sequence | RE[1] | Use | Source |
|---|---|---|---|---|
| pMAD_p1 | CTCCTCCGTAACAAATTGAGG | - | pMAD deletion vector screening and sequencing | M. Atilano |
| pMAD_p2 | CGTCATCTACCTGCCTGGAC | - | | M. Atilano |
| murQ_p1 | TGC**CCATGG**ACAATATGATGCACAATTTATGA | NcoI | Construction of *murQ* deletion construct for insertion into pMAD and deletion of *murQ* from the chromosome of *S. aureus* NCTC8325-4 leaving a 45bp scar | This study |
| murQ_p2 | TCGTTTAACAATGTCACCATTCACTTCTTACACTCCCTAGTTT | - | | This study |
| murQ_p3 | AAACTAGGGAGTGTAAGAAGTGAATGGTGACATTGTTAAACGA | - | | This study |
| murQ_p4 | TGC**AGATCT**CGTTACAATAATATCAATCGCA | BglII | | This study |
| murP_p1 | TGC**CCATGG**CAAGTCCCGTTAGCAGTTCG | NcoI | Construction of *murP* deletion construct for insertion into pMAD and complete deletion of *murP* from the chromosome of *S. aureus* NCTC8325-4 | This study |
| murP_p2 | CATCGCCTTTGTCGTTCCTTAAATCCCTCCTAAGGTTGTCTATC | - | | This study |
| murP_p3 | GATAGACAACCTTAGGAGGGATTTAAGGAACGACAAAGGCGATG | - | | This study |
| murP_p4 | TGC**GGATCC**GCCATTTTCGATTGGGTTAT | BamHI | | This study |
| nagA_p1 | TGC**CCATGG**CACCAGCATCTTGGTTAAAT | NcoI | Construction of *nagA* deletion construct for insertion into pMAD and deletion of *nagA* from the chromosome of *S. aureus* NCTC8325-4 leaving a 15bp scar | This study |
| nagA_p2 | GATTATTTATTAGCTAAATGTGTGGTATAAACCCTTCTTTATTTAT | - | | This study |
| nagA_p3 | ATAAATAAAGAAGGGTTTATACCACACATTTAGCTAATAAATAATC | - | | This study |
| nagA_p4 | TGC**GAATTC**GTAATTCGTCTACATTAAAAGGTT | EcoR1 | | This study |
| murQ_intA | AAGAAGATGACGAGACAAAATATC | - | Checking integration / excision of pMADΔ*murQ,* sequencing of Δ*murQ* deletion region | This study |
| murQ_intB | CACTAAATACGCCACCAATAC | - | | This study |
| murP_intA | GGAGTGTAAGAAGTGATGGAA | - | Checking integration / excision of pMADΔ*murP,* sequencing of Δ*murP* deletion region | This study |
| murP_intB | CAAGCAACTGAATCACATCA | - | | This study |
| nagA_intA | CAGGTTTCTTAGCAGGTTACTT | - | Checking integration / excision of pMADΔ*nagA*, sequencing of Δ*nagA* deletion region | This study |
| nagA_intB | TTCCAGAAGCGTATTCAGTT | - | | This study |
| murQ_SEQ | TGCAAATTGTGAGACAAACG | - | Sequencing of pMADΔ*murQ* and Δ*murQ* deletion region | This study |
| murQ_SEQ2 | ACCTACTGCAGCAATAATTCGTT | - | | This study |
| murP_SEQ | AGGTATCTGAGCATGATGTGAAAG | - | Sequencing of pMADΔ*murQ* and Δ*murP* deletion region | This study |
| nagA_SEQ | ACGGTAGCTTATACAAAGCTGTAA | - | Sequencing of pMADΔ*murQ* and Δ*nagA* deletion region | This study |
| nagA_SEQ2 | ACGAATTACTGCGCCACAT | - | | This study |
| nagA_SEQ3 | CTACTCGTCCATTGATTTTGAA | - | | This study |
| nagA_SEQ4 | CTGATGGTGCACACTTACTTCA | - | | This study |

[1]RE; Restriction enzyme. Added RE sites are shown in bold, and additional sequence to aid in cleavage underlined.
